# Membrane domains and phosphatase exclusion produce robust signaling responses in B cells engaged with natural ligands and artificial cross-linkers presented on lipid bilayers

**DOI:** 10.1101/652719

**Authors:** Marcos Francisco Núñez, Kathleen Wisser, Sarah L Veatch

## Abstract

B cells respond to a wide variety of antigens with varying valiancy and mode of presentation to the highly expressed B cell receptor (BCR). We previously demonstrated that clustering the IgM isotype of BCR with an artificial soluble cross-linker led to the stabilization of an ordered phase-like domain. This domain sorted minimal peptides and full-length proteins to generate local hot-spots for tyrosine phosphorylation at receptor clusters, facilitating receptor activation. BCR can also be activated through interactions with natural ligands or artificial cross-linkers presented at bilayer surfaces, where it is proposed that alternate mechanisms play important roles in receptor clustering and activation, including one mechanism involving the exclusion phosphatases due to their bulky extracellular domains. The goals of the current study are to determine if markers of membrane phases are sorted by BCR clusters formed through engagement with bilayer-presented natural ligands or cross-linkers, and to estimate the contribution that membrane phase partitioning plays in organizing regulatory proteins with respect to BCR clusters under these stimulation conditions. We use super-resolution fluorescence localization microscopy to find that BCR engagement with either a bilayer-presented natural ligand or artificial cross-linker generates ordered phase-like domains that are more robust than those observed with soluble cross-linkers. In addition, we provide evidence that interactions between regulatory proteins and BCR are partially determined through their preference for ordered membrane domains and present a minimal model of receptor activation that incorporates both ordered domains and steric exclusion mechanisms to produce a more sensitive response. Overall, this work highlights that cells are capable of integrating multiple interaction modalities to give rise to cellular functions, likely conferring flexibility and robustness to cellular responses.

## 1 INTRODUCTION

B cells are responsible for reacting to broad stimuli from other cells and within their environment (Youinou, 2007;LeBien and Tedder, 2008;DiLillo et al., 2011). These signals can be initiated through the clustering of cell surface receptors and co-receptors by extracellular ligands and result in cellular-level responses such as the release of cytokines, processing and presentation of antigen peptides to T cells, differentiation, clonal expansion, apoptosis, or combinations of these outcomes (Howard and Paul, 1983;Niiro and Clark, 2002;Monroe and Dorshkind, 2007;Kurosaki et al., 2009). Signaling through the highly expressed B cell receptor (BCR) occurs when the BCR encounters natural ligands, multivalent artificial cross-linkers in solution, or multivalent or monovalent artificial cross-linkers presented on surfaces (Volkmann et al., 2016). One mode of BCR activation involves receptor clustering through binding to ligands or artificial cross-linkers, initiating a signaling cascade that results in phosphorylation of BCR immunoreceptor tyrosine activation motifs (ITAM) by the Src family kinase Lyn (Dal Porto et al., 2004;Harwood and Batista, 2009). Once phosphorylated, the ITAM regions become sites of docking for other signaling mediators, such as Syk, that propagate the cellular-level immune response. Some authors argue that clustering is not required for receptor activation by ligands that engage the antigen binding site of BCRs. Instead, it is proposed that these ligands promote activation through the dissociation of pre-clustered receptors, increasing accessibility of the BCR ITAMs to Syk (Klasener et al., 2014;Gold and Reth, 2019).

While many structural and functional consequences of receptor clustering in B cells are well characterized, mechanisms describing how BCR clustering leads to the robust phosphorylation of BCR ITAMs remains a topic of active debate. One proposed mechanism is that receptor clustering leads to receptor partitioning with ordered membrane domains sometimes referred to as lipid rafts. These domains are enriched in kinase and protected from phosphatase therefore facilitate ITAM phosphorylation and receptor activation. This mechanism is supported by a large quantity of past observations including those using detergent extraction (Yamanashi et al., 1991;Cheng et al., 1999), FRET imaging (Sohn et al., 2006;Sohn et al., 2008), and our recent study that utilized super-resolution fluorescence localization microscopy (Stone et al., 2017a). A second model of receptor activation, referred to as the trans-phosphorylation model, posits that kinases bound to receptors more effectively interact with and activate neighboring receptors when brought into close contact via a multivalent natural ligand or crosslinker (Pierce and Liu, 2010). When natural ligands or artificial cross-linkers are presented on surfaces, additional interactions contribute to receptor activation. Conformational changes within BCR lead to enhanced BCR self-clustering when ligands are presented on surfaces (Tolar et al., 2009;Tolar and Pierce, 2010). Moreover, it is proposed that receptor clusters formed through engagement with surface presented natural ligands or artificial cross-linkers facilitate their own activation by maintaining tight contact between the B cell plasma membrane and the presenting surface, excluding membrane proteins with bulky ectodomains including phosphatases (Davis and van der Merwe, 2006;Depoil et al., 2008;Zhu et al., 2008;Harwood and Batista, 2009). In this model, termed the kinetic segregation model, lowering the concentration of phosphatases proximal to BCRs favors ITAM phosphorylation. Also, it is widely viewed that membrane compartmentalization by the cortical actin cytoskeleton contributes to BCR activation and its coupling to the downstream cellular immune response (Gasparrini et al., 2016;Rey-Suarez et al., 2019). Also, different isotypes of BCR can be compartmentalized within in the plasma membrane, each with distinct local lipid and protein environments that contribute to their activation mechanisms (Klasener et al., 2014;Minguet et al., 2017;Gold and Reth, 2019). It is likely that B cells draw on multiple mechanisms to facilitate antigen recognition, enabling flexibility and robustness for B cells to respond to a diverse repertoire of antigen. In our own past work, we combined essential features of an ordered domain model with the trans-phosphorylation model to produce collective receptor activation within a minimal model of early BCR activation (Stone et al., 2017a). In the current study, we explore the situation where BCR engages with ligands or cross-linkers presented on surfaces where steric phosphatase exclusion is also expected to play an important role.

Plasma membranes isolated from B cells and other cell types separate into coexisting liquid-ordered (Lo) and liquid-disordered (Ld) phases at low temperature (Baumgart et al., 2007;Veatch et al., 2008;Levental and Levental, 2015). Liquid phase domains present in isolated plasma membrane vesicles are macroscopic and have different lipid and protein compositions. In particular, saturated lipids and palmitoylated proteins are enriched in the Lo phase, while unsaturated lipids and single-pass transmembrane proteins are enriched in the Ld phase when they lack palmitoylation sites and are not complexed with other proteins (Diaz-Rohrer et al., 2014;Lorent et al., 2017). Intact B cell plasma membranes do not phase-separate, but instead are predicted to contain domains resembling phases that are small and dynamic and that can be stabilized through interactions with cellular structures such as cortical actin (Veatch et al., 2008;Machta et al., 2011;Zhao et al., 2013). Our past experimental work demonstrated that domains resembling Lo or Ld phases are stabilized in B cell membranes by clustering membrane proteins or peptides with multivalent soluble cross-linkers. Artificially cross-linking the IgM isotype of BCR produces a local environment resembling Lo phase domains. (Stone et al., 2017a). In that case, we found that the sorting of Lyn kinase and CD45 phosphatase based simply on their interactions with membrane domains was sufficient to initiate receptor activation upon clustering with multivalent soluble cross-linkers in a minimal model. A goal of the current study is to identify a role for membrane domains in BCR activation by a natural ligand or by artificial cross-linkers presented on a planar supported bilayer membrane.

In this study, we identify the membrane contribution to B cell signaling in cells engaged with either a bilayer-presented natural monovalent ligand or a bilayer-conjugated artificial cross-linker, and compare to results obtained by clustering BCR with an artificial soluble cross-linker administered in solution. We acquire super-resolution images of fluorescently labeled proteins and peptides alongside BCR on the B cell surface and perform a cross-correlation analysis to quantify the sorting of phase marking peptides and signaling regulators with respect to clusters of BCR. We find that BCR clusters engaged with bilayer-presented natural ligand or artificial cross-linkers robustly sort peptide markers of membrane phases. These phase-like domains, along with additional interactions, determine the local concentration of CD45 and Lyn and likely other signaling molecules involved in initiating a cellular level immune response. We model these interactions and their associated outcomes and obtain qualitative agreement with experimental results and established literature. Overall, we conclude that multiple interaction modalities likely combine to activate BCR and regulate early signaling events when stimulated with surface-presented antigen.

## 2. METHODS

### 2.1 Cell culture

CH27 mouse B-cells (Millipore Cat# SCC115, RRID:CVCL_7178), a lymphoma-derived cell line (Haughton et al., 1986), used in the soluble and surface-presented experiments, were acquired from Neetu Gupta (Cleveland Clinic). Cells were maintained in culture as previously described (Stone and Veatch, 2015). Prior to measurements, cells are typically pelleted from media by spinning 500g for 5 minutes and resuspended into a live cell compatible minimal buffer consisting of (15.0 mM NaCl, 5.0 mM KCl, 1.8 mM CaCl_2_, 1.0 mM MgCl_2_, 6.0 mM Glucose, 20 mM HEPES, pH 7.4).

### 2.2 Large unilamellar vesicle (LUV) formation

Lipids, suspended in chloroform, were dried to form a thin film under N_2_ while being vortexed in a clean glass tube. The lipid film was then placed under vacuum for 30min to remove any residual solvent then hydrated in either purified water or PBS. In some cases, the resulting multilamellar vesicles (MLVs) were subjected to 10 freeze-thaw cycles. LUVs were obtained by extruding the MLVs 30 times with a mini-extruder (Avanti, 610000) using 10mm diameter filters (Whatman Drain Disc, 230300) and a membrane of 0.1μm pore-size (Whatman Nucleopore Track-Etched Membrane, 800309). The final concentration of lipid in LUV suspensions was either 3mg/ml or 1mg/ml. LUVs were stored at 4°C and used within 2 weeks.

### 2.3 Flow chamber and supported planar lipid bilayer assembly

Flow chambers were assembled from a 25×75×1mm plain microscope slide (Fisherbrand, 12-550D) and clean no. 1.5 22×22mm glass coverslip (Fisherbrand, 12-541B) the day of the experiment. Coverslips were cleaned through rinsing with water, ethanol then chloroform before drying under N_2_. In some cases, coverslips were acid cleaned in NoChromix (Sigma-Aldrich Cat# 328693). The microscope slide and coverslip were plasma cleaned for 5 minutes using a plasma cleaner (Harrick Plasma, PDC-32G). 2 strips of PARAFILM were cut into 1 × 2.5 cm and placed on top of the microscope slide. The PARAFILM strips were placed in the middle of the microscope slide with 1 cm of space between them. The coverslip was placed on top of the PARAFILM, and the entire flow chamber was placed on a hot plate at 100°C until the PARAFILM had melted and sealed the slide and coverslip together. PBS was introduced into the chamber to prime.

Planar lipid bilayers were fused from LUVs hydrated in purified water. 50μl of the LUV mixture was diluted 1:1 in PBS then introduced to the flow chamber and incubated at room temperature for 30 minutes, then washed with PBS and incubated for 1 hour at room temperature to allow the bilayer to equilibrate. Prior to the introduction of cells, bilayers are rinsed with the live cell compatible buffer (15.0 mM NaCl, 5.0 mM KCl, 1.8 mM CaCl_2_, 1.0 mM MgCl_2_, 6.0 mM Glucose, 20 mM HEPES, pH 7.4)

For measurements involving bilayer presented streptavidin cross-linkers, 0.5% mol% Biotinyl-DOPE (Avanti, 870273C) is included in the LUV preparation. Following bilayer assembly, membranes are incubated with 200μL of a 1μg/mL streptavidin solution for 10 minutes at room temperature then rinsed in the live cell compatible buffer.

Supported lipid bilayers were also prepared on glass bottomed Mattek petri dishes that were cleaned with ethanol, dried then plasma cleaned prior to incubation with LUV solution diluted in PBS. After incubation, samples were washed taking care that the lipid film was not exposed directly to air.

### 2.4 Calcium mobilization assays

CH27 cells were loaded with 2 μg/mL Fluo-4AM (Invitrogen: F14201) for 5 min at room temperature in 1 mL T/B/S buffer (15.0 mM NaCl, 5.0 mM KCl, 1.8 mM CaCl_2_, 1.0 mM MgCl_2_, 6.0 mM Glucose, 20 mM HEPES, 1mg/ml BSA, 0.25 mM sulfinpyrazone, pH 7.4). The cell suspension was subsequently diluted to a final volume of 15 mL with T/B/S buffer and incubated for 30 min at 37°C to allow for dye loading. After the 30-minute incubation, the cells were spun down at 500g for 5 minutes and re-suspended in 3mL of T/B/S buffer.

For IgM BCR engagement with streptavidin cross-linkers, Fuo-4AM loaded cells were additionally incubated with f(Ab)1-biotin anti mouse IgM (Jackson ImmunoResearch Labs Cat# 115-067-020, RRID:AB_2338587) for 10 minutes at 37°C. Cells were pelleted and resuspended in T/B/S buffer prior to measurements.

For microscopy measurements of calcium mobilization, cells loaded with Fluo-4 were introduced to a Mattek well placed on an inverted microscope stage and imaged over time with 10x magnification. At the early time-points, cells are seen to settle into the focal field, so the number of cells detected varies as a function of time. For soluble stimulation with f(Ab)_2_ αIgM, the soluble antibody was added directly to a Mattek well containing adhered cells. Fluorescent detection of Fluo-4 intensity was accomplished using an incoherent LED light source centered at 488nm (pE excitation system; Cool LED, Andover UK), a quad band filter cube (LF405/488/561/647; Coherent), and an EMCCD camera (iXon-897; Andor, South Windsor, CT). Images were acquired over 10min for each sample with an integration time of 0.5s. Quantification of Fluo-4 intensity within cell boundaries was accomplished from images through an automatic watershed segmentation algorithm in Matlab to obtain the average response over the field of cells. Traces were normalized to the final value for cases where the signal appeared to fall back to the baseline. When possible, traces were normalized to the initial value prior to stimulation. Traces in figures are from single experiments but are representative of results obtained over at least 3 independent measurements.

For measurements of calcium mobilization in solution, 100,000 cells loaded with Fluo-4 were placed per well in a 96-well plate and imaged a microplate plate reader (SpectraMax iD33; Molecular Devices, San Jose, California). 25μL of LUVs were added to 175μL of cells at time = 0 such that the final concentration of lipid was 12.5μg/ml. Temperature was held at 30°C for all samples to ensure the POPE LUVs would be in a liquid state. Fluorescence intensity was measured at 0.5s increments for 10min. Raw fluorescence intensity was normalized by the values detected <5sec after the injection, before the detected rise in fluorescence emission intensity. Traces shown represent the average values of normalized curves over at least 3 independent measurements.

### 2.5 Fluorophore and biotin conjugation to secondary antibodies

Biotinylation and Atto555 conjugation of Goat f(Ab)_1_ anti-Mouse IgM (Jackson ImmunoResearch Labs Cat# 115-007-020, RRID:AB_2338477) was accomplished simultaneously. 2.25μL of 15 mM amine reactive biotin-X, SE (Invitrogen: B1582) and 0.45μL of 10mM NHS-Ester Atto655 (Millipore-Sigma: 76245) were mixed with 150μL of 1.3mg/mL of the f(Ab)_1_ fragment at pH 8.5 and rotated for 1 hour at room temperature in an aqueous solution buffered by 0.01M NaH_2_PO_4_ with 0.01M NaH_2_CO_3_. After incubation, the modified f(Ab)_1_ fragment was purified and separated from unbound dye through a gel filtration column (GE Healthcare illustra NAP Columns from Fisher: 45-000-151) in 1X-PBS+1mM-EDTA. The modified antibody was further purified by centrifugation in a Vivaspin-500 Polyethersulfone concentration spin column with a 30kDa cutoff (Vivaspin: VS0121). The modified antibody was then conjugated to additional NHS-Ester Atto655 a second time by mixing 0.4μL of 10mM of the NHS-Ester Atto655 with the modified antibody at room temperature in pH 8.5 for 1 hour. The antibody was purified through the gel column and spin column again before its optical spectrum was obtained to estimate degree of label (~3 dye/antibody).

Biotinylation and silicon rhodamine (SiR) conjugation of Goat f(Ab)_1_ anti-Mouse IgM was accomplished similar to the above.3.5μL of 15 mM amine reactive biotin-X, SE (Invitrogen: B1582) and 2.0μL of 10mM NHS-Ester SiR (Spirochrome: SC003) were mixed with 300μL of 1.3mg/mL of the f(Ab)_1_ fragment at pH 8.5 and rotated for 1 hour at room temperature in an aqueous solution buffered by 0.01M NaH_2_PO_4_ with 0.01M NaH_2_CO_3_. After incubation, the modified f(Ab)_1_ fragment was purified as described above.

In some cases, secondary antibodies were fluorescently labeled following similar procedures as outlined above, but the amine reactive biotin-X was left out of the reaction mixture.

### 2.5 DNA constructs and transient transfection

The plasmids used for this study were used and described previously (Stone et al., 2017a). Briefly, Lyn-eGFP and PM-eGFP plasmids (Pyenta et al., 2001) were gifts from Barbra Baird and David Holowka (Cornell University, Ithaca, NY) and were cloned using standard techniques to replace eGFP with mEos3.2. Plasmid DNA encoding mEos39.2 protein and YFP-TM were gifts from Akira Ono (University of Michigan, Ann Arbor, MI). The YPF-TM plasmid encodes the transmembrane domain of LAT (Linker for Activation of T-cells) fused to the YFP such that the fluorescent protein is on the extracellular side of the cell. This plasmid was cloned using standard techniques to replace YFP with mEos3.2. The mEos3.2 tagged constructs used here are in Clontech N1 plasmid vector background (Clontech, Mountain View, CA).

CH27 cells were transiently transfected by Lonza Nucleofector electroporation (Lonza, Basel, Switzerland) with electroporation program CA-137. Typically, 10^6^ CH27 cells were transfected with 1.0 mg of plasmid DNA. For soluble measurements, transfected cells were grown on glass-bottom Mattek petri dishes overnight for next-day sample preparation. For surface-presented measurements, transfected cells were grown overnight in T-25 cell culture flask with filter caps in 5mL of growth media. The following day, the transfected cells were harvested and pelleted at 500g for 5 minutes before being labeled for sample preparation.

### 2.5 Preparation of samples for fluorescence microscopy

For stimulation of CH27 cells by PC membranes, unlabeled live CH27 cells were suspended in a live cell compatible minimal buffer at room temperature (15.0 mM NaCl, 5.0 mM KCl, 1.8 mM CaCl_2_, 1.0 mM MgCl_2_, 6.0 mM Glucose, 20 mM HEPES, pH 7.4) then introduced into a flow chamber presenting a PC containing membrane. Cells were incubated for 5 min prior to chemical fixation in 2% paraformaldehyde (PFA), 0.15% glutaraldehyde in 0.5X PBS for 20min. Fixation was quenched by incubating with PBS containing 3% Bovine serum albumin (BSA) for 20 minutes at room temperature.

For stimulation of CH27 cells by surface-presented streptavidin, live CH27 cells in suspension were pelleted at 500g for 5 minutes at 4°C then incubated on ice for 10 minutes with 5μg/mL of anti IgM f(Ab)_1_ modified with biotin and Atto655 in the live cell compatible buffer defined above. Cells were washed twice through centrifugation at 500g for 5 minutes at 4°C and ultimately suspended in room temperature Tyrode’s buffer. Cells were then introduced into the flow chamber containing a streptavidin-loaded bilayer and incubated for 5 minutes in Tyrode’s buffer at room temperature prior to chemical fixation.

For BCR labeling and stimulation using soluble streptavidin, CH27 cells were incubated in a Mattek well overnight to facilitate their natural adhesion to glass. Cells were then labeled in 5μg/mL of anti IgM f(Ab)_1_ modified with biotin and Atto655 in the live cell compatible buffer for 10 min at room temperature. Cells were rinsed in the live cell compatible buffer before incubation with streptavidin (1.0μg/mL) for 5 min followed by chemical fixation.

For cells engaged with surface-bound transferrin, 2mg/ml biotinylated BSA was incubated with a clean flow chamber for 10 min at room temperature. After washing, flow cells were incubated with 1μg/mL streptavidin solution for 10 minutes at room temperature. Live CH27 cells in suspension were pelleted at 500g for 5 minutes at 4°C then incubated on ice for 10 minutes with 3.5μg/mL biotinylated transfer-rin (Sigma-Aldrich: T3915) in the live cell compatible buffer. Cells were washed twice through centrifugation at 500g for 5 minutes at 4°C and ultimately suspended in room temperature Tyrode’s buffer before loading into the flow chamber where they were allowed to adhere for 5min prior to chemical fixation.

For phosphorylated BCR (pBCR) detection, chemically fixed cells were permeabilized in 0.1% Triton-X 100 then labeled with anti-human phosphotyrosine CD79A (Tyr182) rabbit antibody (Cell Signaling Technology Cat# 5173, RRID:AB_10694763). The primary antibody was detected using one of two a secondary goat anti rabbit polyclonal antibodies. A commercially labeled Alexa 532 secondary was used (Invitrogen: A-11009) for diffraction limited images and for images acquired alongside IgM. For super-resolution measurements alongside transiently expressed peptides, an unlabeled secondary antibody (Jackson ImmunoResearch Labs Cat# 111-005-003, RRID:AB_2337913) was conjugated to Atto655 using methods described above.

For PhosphoTyrosine (pTyr) detection, chemically fixed and permeabilized cells were labeled with anti-phosphotyrosine 4G10 Platinum mouse monoclonal antibody of the IgG2b subclass (Millipore Cat# 05-1050, RRID:AB_916371). The primary antibody was detected using a secondary goat anti mouse IgG Fcγ subclass 2b antibody (Jackson ImmunoResearch Labs Cat# 115-005-207, RRID:AB_2338463). The secondary antibody was conjugated to Alexa Fluor 532 using methods described above.

For CD45 detection, chemically fixed CH27 B cells were incubated with anti-mouse CD45R (B220) primary antibody clone RA3-6B2 conjugated directly to Alexa 532 (Thermo Fisher Scientific Cat# 14-0452-81, RRID:AB_467253).

### 2.6 Imaging

Imaging was performed using an Olympus IX81-XDC inverted microscope. TIRF laser angles where achieved using a cellTIRF module, a 100X UAPO TIRF objective (NA = 1.49), and active Z-drift correction (ZDC) (Olympus America) as described previously (Stone and Veatch, 2014;2015;Stone et al., 2017a). Atto 655 and SiR were excited using a 647 nm solid state laser (OBIS, 100 mW, Coherent) and Alexa 532 was excited using a 532 nm diode-pumped solid-state laser (Samba 532–150 CW, Cobolt). Photoactivation of mEos3.2 was accomplished with a 405 nm diode laser (CUBE 405-50FP, Coherent) and simultaneously excited with a 561 nm solid state laser (Sapphire 561 LP, Coherent). Simultaneous imaging of Atto655 or SiR and mEos3.2 was accomplished using a LF405/488/561/647 quadband filter cube (TRF89902, Chroma, Bellows Falls, VT), while imaging of Atto655 or Sir and Alexa532 accomplished using a 405/488/532/647 quadband filter cube (Chroma, Bellows Falls, VT). Emission was split into two channels using a DV2 emission splitting system (Photometrics) with a T640lpxr dichroic mirror to separate emission: ET605/52m to filter near-red emission, and ET700/75m to filter far-red emission (Chroma).

Diffraction limited images were acquired by maintaining laser intensities and TIR illumination angles to enable comparison of intensities observed across samples. Cells within individual images were masked by hand, and pixel values within the mask were used to obtain average values over the cell footprint or used individually to generate histograms of pixel intensities.

Interference Contrast Microscopy (ICM) images were acquired by illuminating samples with a broad band incoherent light source centered at 565nm (pE excitation system; Cool LED, Andover UK). Collected light was filtered using a U-C QUAD T2 cube (TIRF C155405, Chroma, Bellows Falls, VT).

Super-resolution localization microscopy images were obtained by adjusting laser intensities such that single fluorophores could be distinguished in individual frames, and were generally between 5 kW/cm^2^ and 20 kW/cm^2^. Integration times were maintained at 20ms and at least 10,000 individual images were acquired per cell. Samples labeled with BCR-Atto 655 and mEos3.2 were imaged in a buffer found to be optimal for Fluorescence Localization Microscopy with these probes: 30 mM Tris, 9 mg/ml glucose, 100 mM NaCl, 5 mM KCl, 1 mM KCl, 1 mM MgCl_2_, 1.8 mM CaCl_2_, 10 mM glutathione, 8 mg/ml catalase, 100 mg/ml glucose oxidase, pH 8.5 (Stone et al., 2017a). Samples labeled with Atto 655 and Alexa 532 were imaged in a buffer more suitable for the inorganic dye pair (Heilemann et al., 2009;Stone et al., 2017a): 50 mM Tris, 100 mg/mL glucose, 10 mM NaCl, 100 mM 2-mercaptoethanol, 50 mg/ml glucose oxidase, 200 mg/ml catalase, pH 8. Samples labeled with SiR and Alexa532 were imaged in: 50 mM Tris, 100 mg/mL glucose, 10 mM NaCl, 100 mM 2-mercaptoethanol, 10 mg/ml glucose oxidase, 200 mg/ml catalase pH 8.5.

### 2.7 Single molecule localization and super resolution image reconstruction

Candidate finding for single-molecule fitting was accomplished by fitting the intensity profile of local maxima in background subtracted and wavelet filtered images using 2-D Gaussian functions as described previously (Stone et al., 2017a). Once candidates were fit and localized, they were culled to remove outliers based on size, intensity, and localization error for each color using software described previously (Veatch et al., 2012). The near-red channel was overlaid onto the far-red channel using an image transform matrix calculated from fiducial bead marker images obtained during each experiment. The images were aligned to account for stage drift which resulted in the final reconstructed STORM image from the culled, transformed, and aligned localizations. Final image resolution and the surface density of probes were calculated for each color using previously described methods (Veatch et al., 2012).

Localizations used to generate reconstructed images are grouped such that probes detected in sequential image frames within a specified radius are averaged to single localizations. Reconstructed images shown in figures are generated from a 2D histogram of grouped localizations with pixels of 25nm. In main images, histograms are filtered with a 2D Gaussian function with standard deviation of 50nm, which is greater than the resolution of the image, which is typically 20nm-30nm. Insets within images are histogram images drawn without filtering.

### 2.8 Correlation analysis

Auto-correlation and Cross-correlation functions were tabulated from masked, reconstructed super-resolution images using previously described methods (Sengupta et al., 2011;Veatch et al., 2012;Stone and Veatch, 2015;Stone et al., 2017a). Images are reconstructed with 25nm pixels from ungrouped localizations for cross-correlations and from grouped images for auto-correlations and in both cases, images are not further blurred prior to tabulating correlation functions. Average cross-correlation functions are obtained through a simple average over single cell measurements, and errors presented on average curves represent the standard error of the mean.

### 2.9 Simulations of receptor activation

A conserved order parameter 2D Ising model was simulated on a 256 by 256 square lattice as described previously (Machta et al., 2011;Burns et al., 2016;Stone et al., 2017a) with minor modifications. Briefly, components that prefer ordered or disordered regions are represented as pixels that have value of S = +1 and S = −1 respectively, and an equal number of +1 and −1 components were included in all simulations. The vast majority of +1 and −1 pixels represent unspecified membrane components representing proteins and lipids. In addition, 200 pixels with values of +1 are classified as receptors, 100 pixels with values +1 are classified as kinases, and 100 pixels with values −1 are classified as phosphatases. Receptors are clustered by applying a strong attractive circular field (φ^R^) at the center of the simulation frame that only acts on receptors. In some simulations, a repulsive field with the same dimensions as φ^R^ is applied that only acts on phosphatases (φ^P^). The final Hamiltonian is given by:

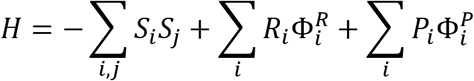

The first term sums over the four nearest neighbors (j) surrounding the pixel i and applies to all components. The second term only contributes when receptors occupy position i, where R_i_=1, otherwise R_i_=0. The Third term only contributes when phosphatases occupy position i, where P_i_=1, otherwise P_i_=0. The fields Φ^R^i and Φ^R^i have a circular shape with a radius of 48 pixels (~100 nm) and is centered in a simulation box with periodic boundary conditions.

Components are moved around the simulation box using a Monte Carlo algorithm that maintains detailed balance. Energy is calculated for proposed moves and moves are accepted stochastically with probability exp(−βΔH) where β is the inverse temperature (T) multiplied by Boltzmann’s constant (k_B_) and ΔH is the change in energy between initial and final states. Simulations were run at either 1.1T_C_, for models with membrane domains, or 10T_C_ for models without membrane domains, where T_C_ = 2/ln(1+sqrt(2)) is the critical temperature of the 2D Ising model. The receptor and phosphatase fields are made independent of the temperature of the membrane by multiplying by T/T_C_ so that the acceptance probability of moves in and out of the field does not vary with β. The magnitude of the receptor field is varied between −.01 and −100 k_B_T/T_C_ filling the range of weak to complete receptor clustering. The magnitude of the phosphatase field was held constant at +10 k_B_T/T_C_, a value that ensured complete phosphatase depletion.

If a move is accepted that places a receptor neighboring a kinase, then the receptor is phosphorylated at a low probability (0.15%). If a move is accepted that places a receptor neighboring a phosphatase, then the receptor is dephosphorylated at a high probability (100%). These probabilities are chosen to produce a low level of phosphorylation in simulations that contain an equal number of kinases and phosphatases with unclustered receptors and do not satisfy detailed balance. When receptors are activated they gain kinase activity, in that a move that places an activated receptor next to a second receptor results in the second receptor becoming phosphorylated at a low probability (0.15%).

All simulations were initially run using non-local exchanges to decrease equilibration times. After equilibration, exchanges were then restricted to nearest neighbors in order to mimic diffusive dynamics. Simulation sweeps were converted to time assuming a diffusion coefficient of roughly 4 μm2/s, with one sweep corresponding to roughly 1 μs. Simulations were recorded for 1000 sweeps which corresponds to roughly 1 s.

All analyses were carried out in MATLAB (MATLAB, RRID:SCR_001622).

## 3 RESULTS

### 3.1 PC lipids incorporated into supported lipid bilayers are a natural ligand for CH27 B cells

The CH27 cells used in this study express an IgM isotype of the BCR, and past work indicates that this IgM is activated upon specific engagement with phosphatidyl choline (PC) lipids adsorbed on soluble proteins, intact red blood cells, or liposomes (Haughton et al., 1986;Mercolino et al., 1986). To verify this, we probed calcium mobilization and stained for phosphorylated BCR in CH27 cells exposed to PC lipids and cross-linkers that cluster and activate B cells without engaging the natural antigen binding site of the receptor. We probed calcium mobilization in Fluo-4 loaded CH27 cells using two assays. First, cells were deposited onto a glass bottom dish with a planar supported lipid bilayer (PLB) and allowed to settle over time while visualizing Fluo-4 emission intensity as a proxy for cytoplasmic calcium concentration using fluorescence microscopy (Fig 1A). An increase in cytoplasmic calcium concentration was observed when cells settled onto a supported lipid bilayer of PC lipids or when cells pre-labeled with biotinylated f(Ab)_1_ against IgMμ (αIgM) settled on a PC bilayer presenting streptavidin cross-linkers. Calcium mobilization was not observed when cells settled on membranes made exclusively of phosphatidyl ethanolamine (PE) lipids, or when cells settle directly onto a clean glass coverslip. In these experiments, cells settle on the dish over time, broadening their response. In addition, curves sometimes start below 1 because cells appear dimmer as they come into focus. For comparison, CH27 cells naturally adhered to a glass coverslip were also stimulated with a soluble f(Ab)_2_ αIgM cross-linker. In this control, cells were exposed to cross-linkers simultaneously and exhibit a sharp and collective response. We also probed the calcium response for cells incubated with 100nm diameter large unilamellar vesicles (LUVs) in suspension via a plate reader fluorimeter assay (Fig 1B). Here we found that calcium mobilization occurs when CH27 cells are incubated with POPC LUVs, but not appreciably when cells are incubated with POPE LUVs.

**Figure 1:**
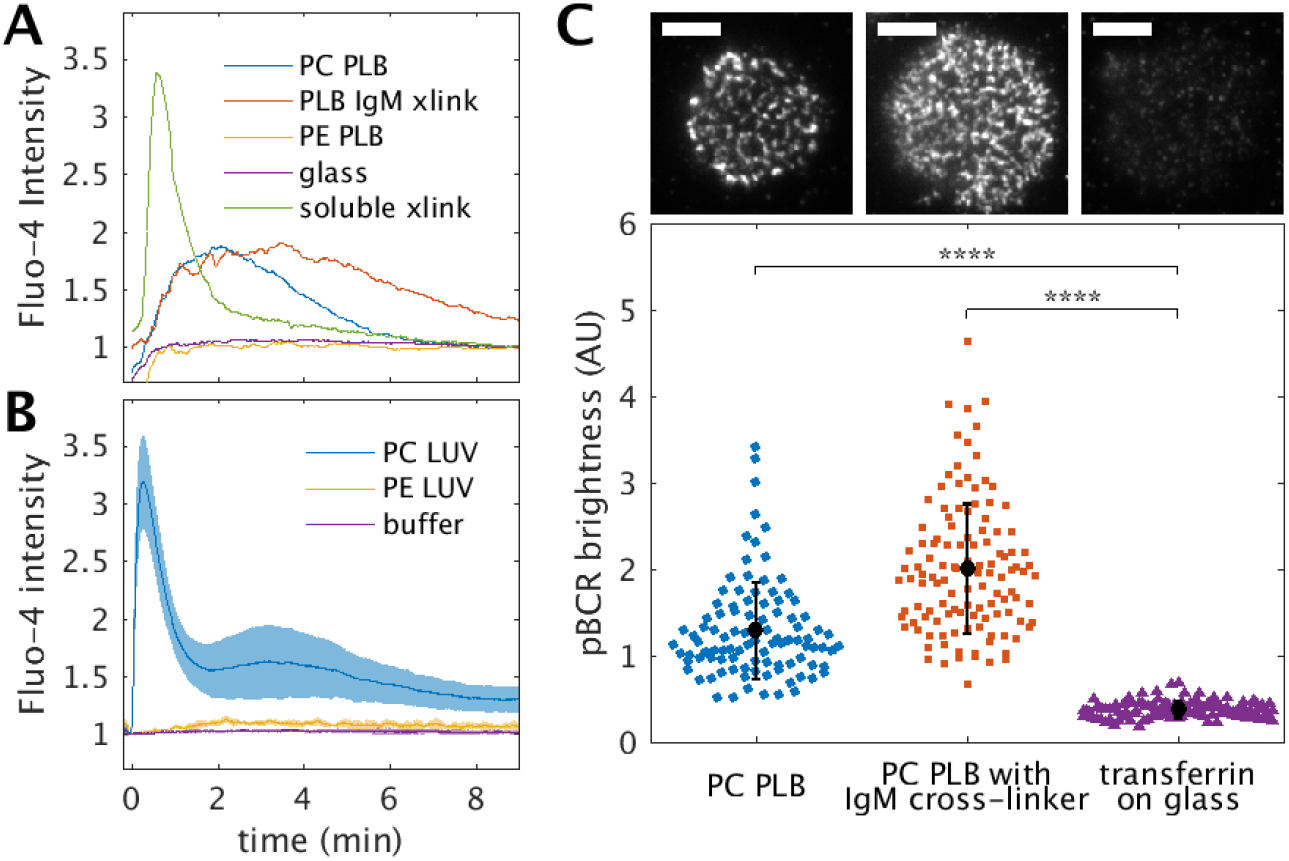
CH27 B cells robustly activate through their BCR upon engagement with PC lipid containing membranes. (A) Calcium mobilization is detected using the cytoplasmic calcium indicator Fluo-4 in CH27 cells deposited onto planar lipid bilayers (PLBs) containing PC lipids. Here the average Fluo-4 level across a field of cells is plotted vs. time after cells are introduced into the chamber. Calcium also mobilizes when biotin-f(Ab)_1_ αIgM labeled cells are exposed to bilayer presented or soluble streptavidin as an artificial cross-linker. (B) Calcium mobilization is also detected in CH27 cells probed in suspension upon exposure large unilamellar vesicles (LUVs) of POPC but not appreciably in cells exposed to LUVs of POPE. (C) pBCR staining of CH27 cells imaged under total internal reflection indicates that BCR is phosphorylated in cells engaged with PC membranes (top left) or when IgM BCR is artificially clustered with bilayer presented streptavidin (top middle) compared to an unstimulated control conjugated to a glass surface through biotinylated transferrin (top right). Images are acquired under the same illumination conditions and displayed the same intensity scale. All scale-bars are 5μm. The quantification (bottom) shows the average brightness across the cell footprint over <100 individual cells for the three conditions shown.

In order to verify that calcium mobilization was a consequence of signaling through the BCR, we stained for activated BCR using an antibody against CD79A (Igα) phosphorylated at Tyr182 (pBCR). CH27 cells chemically fixed and stained 5 min after coming into contact with a PC lipid bilayer exhibit punctate pBCR staining when imaged in total internal reflection (TIR), indicating that a pool of BCR is activated upon engagement with the PC lipid membrane (Fig 1C). pBCR staining is also evident in CH27 cells labeled with biotinylated f(Ab)_1_ αIgM and cross-linked with streptavidin presented on a supported bilayer, but not in control cells which are adhered via biotinylated transferrin to glass pre senting streptavidin. pBCR staining indicates that activated BCRs are organized to extended, sometimes non-circular domains at the cell surface, in agreement with previous studies that activate BCR using bilayers conjugated to a range of natural ligands and artificial cross-linkers (Carrasco et al., 2004;Carrasco and Batista, 2006a;Fleire et al., 2006). pBCR staining in PC lipid-engaged cells is organized into tighter clusters than pBCR staining in streptavidin engaged cells. Fig 1D shows a quantification of the average pBCR brightness across multiple cells, demonstrating that both bilayers presenting PC lipids and artificial IgM cross-linkers activate BCR to similar levels. Together, these measurements support past observations that CH27 cells are activated specifically by PC lipid headgroups and that activation occurs through engagement of BCR. Notably, since individual PC lipids contain a single PC chemical group, they bind the receptor with monomeric valency.

### 3.2 Markers of ordered and disordered phases sort with respect to BCR clusters engaged with PC lipid membranes

We next monitored the sorting of two mEos3.2 conjugated minimal peptides with respect to clusters of phosphorylated BCR in CH27 cells in contact with PC lipid bilayers using super-resolution fluorescence localization microscopy. In past work, we demonstrated that minimal membrane anchored peptides differentially sort with respect to phase separated liquid-ordered (Lo) and liquid-disordered (Ld) domains in isolated giant plasma membrane vesicles (GPMVs) and BCR clusters in intact CH27 B cells (Stone et al., 2017a). The PM peptide is the palmitoylated and myristoylated anchor sequence from that partition with the Lo phase in GPMVs and with BCR clusters in cells (Pyenta et al., 2001;Stone et al., 2017a). The TM peptide is the transmembrane helix from LAT with mutations at palmitoylated cysteines that partitions with the Ld phase in GPMVs and away from BCR clusters in cells (Levental et al., 2010;Lorent et al., 2017;Stone et al., 2017a). In this past work, soluble streptavidin was used as an artificial cross-linker for BCRs labeled with biotinilated f(Ab)_1_ αIgM. Here we ask if a similar trend is observed when BCR is engaged with a bilayer-presented natural ligand.

CH27 cells transiently expressing either PM or TM peptides conjugated to the photo-activatable probe mEos3.2 were introduced onto glass-supported PC planar bilayers within simple flow chambers at room temperature as described in Materials and Methods. The use of a flow chamber reduces the time required for cells to settle onto the bilayer surface, enabling us to probe the relatively short activation time (5 min) for comparison with previous work. After incubating on the bilayer for 5 minutes, cells were chemically fixed, then pBCR was labeled with primary antibody, and visualized with an Atto655 conjugated secondary antibody. Super-resolution imaging was accomplished using fluorescence localization microscopy by detecting mEos3.2 and Atto655 signals simultaneously under conditions that allowed for stochastic activation. Single molecules localized within at least 10,000 acquired images were used to reconstruct the representative super-resolution images shown in Fig 2A.

**Figure 2:**
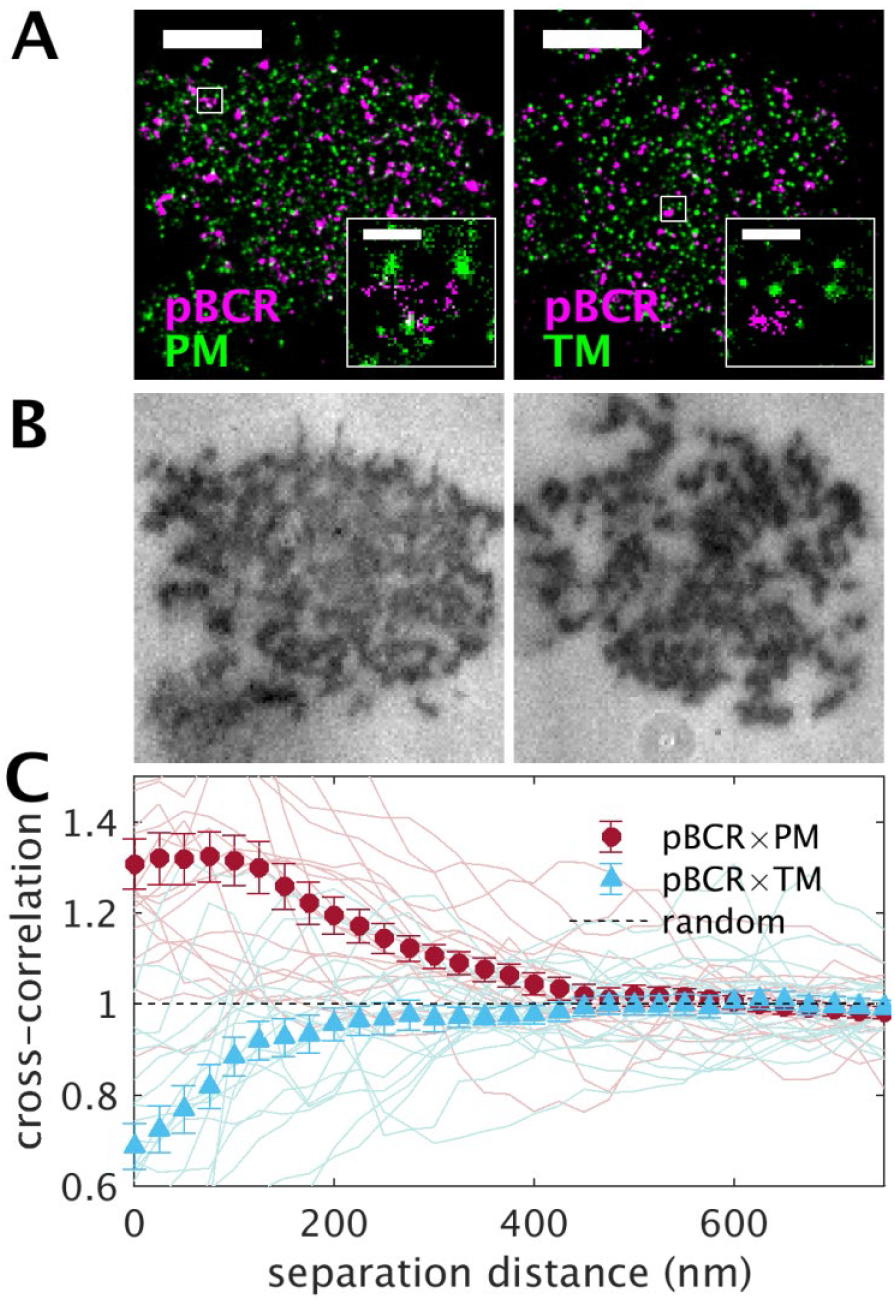
Phase-marking minimal peptides sort with respect to BCR clusters formed through engagement supported bilayers of DOPC. (A) Representative re-constructed super-resolution images of pBCR antibody stained with Atto655 and the minimal membrane anchored peptides PM (left) or TM (right) conjugated to mEos3.2. Scale bars are 5μm in main images and 500nm in insets. (B) ICM images report on the distance between the plasma membrane and the glass support and indicate that these cells exhibit significant membrane topography across the basal cell surface. (C) Cross-correlation functions tabulated for 20 cells between pBCR and PM or 20 cells pBCR and TM. Curves from individual cells are shown as lighter color lines and average curves are shown as filled symbols with error bounds representing the standard error of the mean.

Similar to the diffraction limited images of Fig 1C, pBCR staining in super-resolved images indicate activated BCRs are organized into extended clusters at the cell surface. In contrast, both PM and TM peptides appear as small puncta that are distributed more evenly across the cell surface. These tight puncta are most likely a consequence of observing the same mEos3.2 fluorophore multiple times over the course of imaging. This over-counting effect is common to super-resolution localization measurements and does not indicate that peptides are themselves organized into clusters (Veatch et al., 2012;Stone et al., 2017b). Cells were also imaged using interference contrast microscopy (ICM; Fig 2B). In these images, contrast arises from destructive interference between light reflected off the coverslip and membrane, with darker pixels indicating regions of closer contact (Weber, 2003). CH27 cells engaged with PC bilayers exhibit significant surface topography on their basal surface, with pBCR staining occurring largely at positions where the cell and supported membrane are in close contact.

Co-distributions of pBCR with minimal peptides were quantified using pair cross-correlation functions, g(r), tabulated from masked regions of cells as described in Materials and Methods. The cross-correlation function reports the relative density of localizations of one color probe a distance r from the average probe of the other color within a specified region of interest. g(r) is normalized so that values greater than 1 indicate co-clustering, values less than 1 indicate depletion, and values equal to 1 indicates a random co-distribution within the specified error bounds. Fig 2C shows curves tabulated for individual cells along with the average curve for each peptide. Error bounds on average curves represent the standard error of the mean between cells. One advantage of this quantification method is that it is largely independent of the density of probes present, beyond effects on signal to noise. We detect no appreciable correlation between the amplitude of g(r) and the expression level of transiently expressed peptides in these measurements (Supplementary Figure 1).

The cross-correlation functions in Fig 2C indicate that PM is robustly enriched and TM is robustly depleted from pBCR labeled clusters engaged with PC PLBs. The average cross-correlation curve between pBCR and PM has a maximum of g(r) = 1.31±0.04 for r<25nm, indicating that the average concentration of PM is roughly 30% higher within pBCR labeled clusters than in the membrane overall. In contrast, the average cross-correlation curve between BCR and TM has a minimum of g(r) = 0.70 ±0.04 for r<25nm, indicating that the concentration of TM is 30% lower in pBCR labeled clusters than in the membrane as a whole. Interestingly, the range of PM enrichment is larger than the range of TM depletion, which is roughly equivalent to the size of the BCR clusters themselves (Supplementary Figure 2). Past work has shown that these probes are randomly distributed with respect to IgM labeled BCR in the absence of stimulating ligands or cross-linkers, within experimental limitations and signal to noise (Stone et al., 2017a). The minimal peptides PM and TM are not known to exhibit direct interactions with proteins, therefore we conclude that their sorting behavior arises from their interactions with membranes. Peptide sorting with respect to these BCR clusters mimics the sorting observed with respect to liquid-ordered phase domains in isolated vesicles and we refer to these clusters as ordered phase-like domains.

### 3.3 Markers of ordered and disordered phases sort with respect to BCR clusters engaged with a bilayer-presented cross-linker

The majority of past work uses artificial, antibody based cross-linkers to cluster and activate BCR, although evidence suggests that activation strategies wherein epitope interaction occurs through the antigen binding site and those that bypass the binding site through cross-linking can occur via alternate mechanisms (Pierce and Liu, 2010;Mukherjee et al., 2013) To address this, we also probed the activation of BCR in CH27 cells in which biotinylated and fluorescently labeled f(Ab)_1_ antibodies bound to BCR IgM were cross-linked with streptavidin presented on a planar supported bilayer. αIgM f(Ab)_1_ labeled cells were introduced into a flow chamber containing a streptavidin-loaded supported DOPC lipid bilayer at room temperature and stimulated for 5min. Separate measurements using fluorescently labeled streptavidin verified that the cross-linker was mobile (Supplementary Figure 3). Cells were then chemically fixed and stained for pBCR, and imaged in TIR using diffraction limited (Fig 3A) or super-resolution localization (Fig 3B) microscopy. IgM staining is visually co-localized with pBCR in streptavidin-engaged cells, with pBCR labels decorating connected f(Ab)_1_ αIgM labeled clusters. This indicates that subsets of αIgM-labeled receptors are also tagged with the pBCR label. The co-localization of these two probes is quantified by comparing the cross-correlation function of pBCR and IgM with the autocorrelation of IgM (Fig 3C). The auto-correlation is similar to the cross-correlation, but reports the relative density of localizations of one color probe a distance r from the average probe of the same color within a specified region of interest. The pBCR×IgM cross-correlation and IgM autocorrelation functions are similar in shape and magnitude, with the IgM autocorrelation extending to slightly higher values than the cross-correlation at short distances. This difference most likely arises from multiple observations of single IgM molecules within images, which contribute to autocorrelation functions but not to cross-correlations between distinct labels (Veatch et al., 2012). It is possible that some BCR labeled by f(Ab)_1_ αIgM become activated through engagement with PC lipids rather than via the streptavidin cross-linker, but due to technical limitations we were unable to conduct these measurements on supported lipid bilayers that lacked PC headgroups (Sendecki et al., 2017). Importantly, there is no significant pool of pBCR not also marked by the αIgM probe which would indicate that BCR becomes activated by the PC bilayer away from cross-linked αIgM clusters.

**Figure 3:**
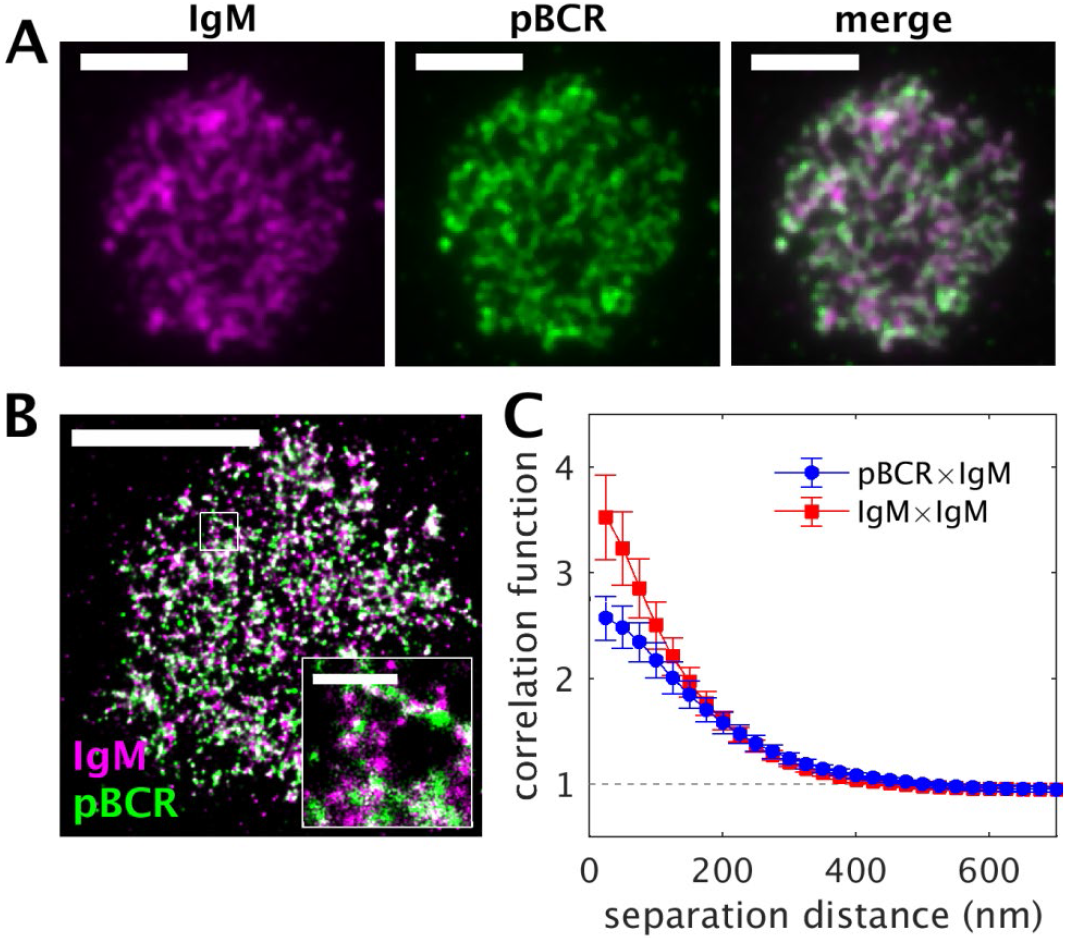
IgM labels reflect lateral organization of activated BCR in CH27 cells engaged with an artificial cross-linker presented on a DOPC membrane. (A) Diffraction limited images of biotin-f(Ab)_1_ αIgM and pBCR indicate significant co-localization of these two probes. Scale bars are 5μm. (B) Colocalization is also apparent a representative super-resolved image of a different cell labeled with the same probes. Scale bar is 5μm in main image and 500nm in inset. (C) The average cross-correlation function between IgM and pBCR has roughly the same amplitude and shape as the auto-correlation function for IgM (9 cells), indicating that both probes exhibit similar spatial distributions.

Figure 4 shows representative images and quantifications of biotinylated αIgM and phase-marking peptides in CH27 cells engaged with PC bilayer presented streptavidin. Similar to the images of Fig 2, BCR organizes into extended and often elongated clusters while TM or PM peptides more randomly sample the cell surface. Different from the images of pBCR in Fig 2, αIgM labeled BCR occupies a larger fraction of the cell surface, sometimes forming connected structures. Also, ICM images of IgM conjugated cells exhibit similar contrast across the cell surface, indicating reduced membrane topography compared to PC conjugated cells. Subtle enrichment of PM and especially exclusion of TM from IgM BCR clusters is visually evident in reconstructed super-resolution images cells engaged to the bilayer through the artificial cross-linker streptavidin (insets in Fig 4A and additional cells in Supplemental Figures 4,5).

**Figure 4:**
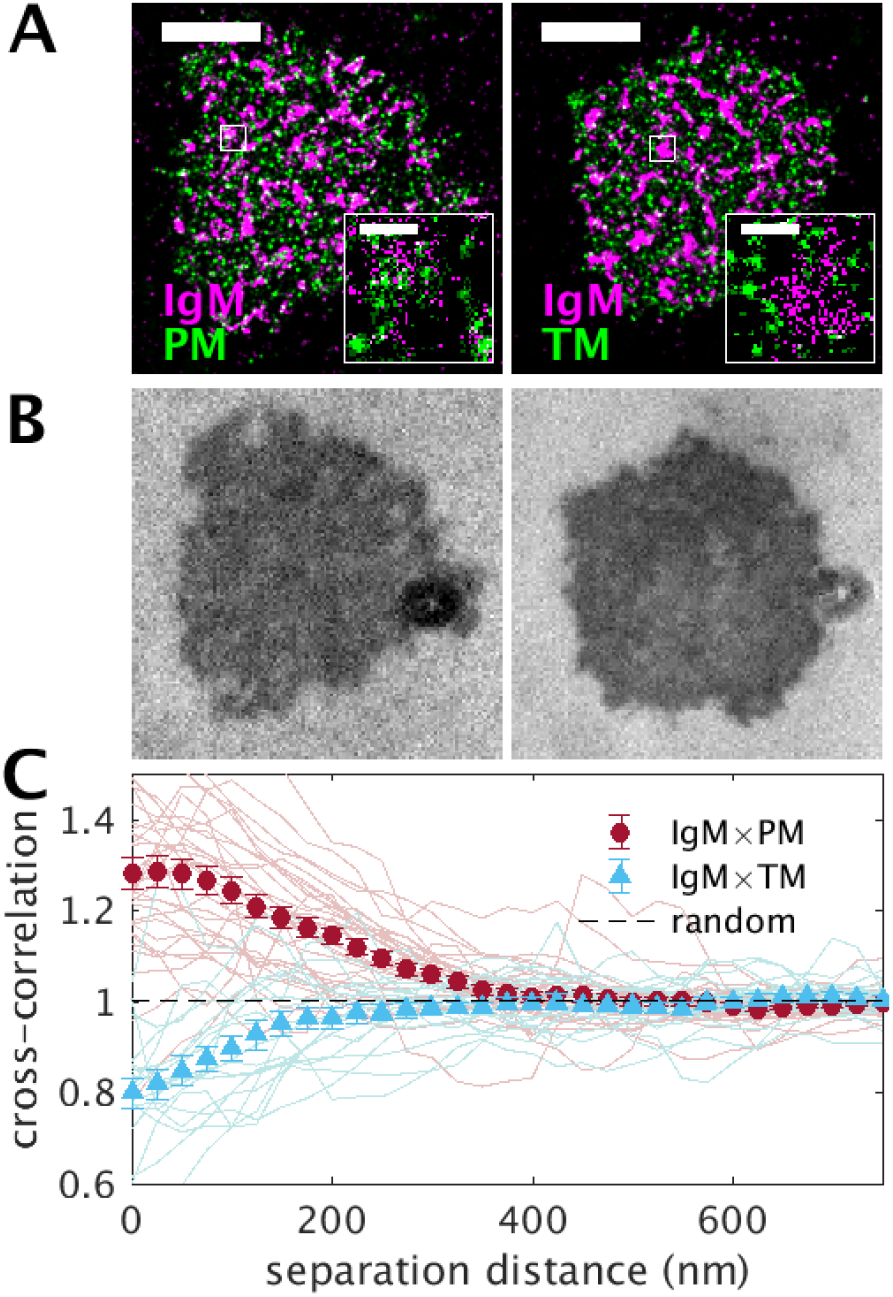
Phase-marking minimal peptides sort with respect to BCR clusters formed through engagement with POPC membranes presenting the artificial cross-linker streptavidin. (A) Representative reconstructed super-resolution images of BCR clusters, marked with biotinylated f(Ab)_1_ αIgM conjugated to Atto655 and the minimal membrane anchored peptides PM (right) or TM (left) conjugated to mEos3.2. Scale bars are 5μm in main images and 500nm in insets. (B) ICM images report on the distance between the plasma membrane and the glass support and indicate that these cells exhibit minimal membrane topography across the basal cell surface. (C) Cross-correlation functions tabulated for 31 cells between IgM and PM or 19 cells IgM and TM. Curves from individual cells are shown as lighter color lines and average curves are shown as filled symbols with error bounds representing the standard error of the mean between cells.

Cross-correlation functions tabulated from individual cells and their averaged curves are shown in Fig 4C and indicate that PM is preferentially enriched while TM is depleted from BCR clusters. Again, we observe no dependence on the expression level of peptide probes (Supplemental Figure 1). The average cross-correlation curve between BCR and PM has a maximum of g(r)=1.28±0.04 for r<25nm, indicating that the average concentration of PM is roughly 30% higher within BCR clusters than in the membrane overall. The average cross-correlation curve between BCR and TM has a minimum of g(r)=0.80 ±0.03 for r<25nm, indicating that the concentration of TM is 30% lower in BCR clusters than in the membrane as a whole. Similar to observations of Fig 2C for cells engaged with PC bilayers, we observe that the range of PM enrichment is larger than the range of TM depletion when BCR engages with a bilayer presented cross-linker (Supplementary Figure 2). Again, we use these findings to conclude that BCR clusters engaged with an artificial, bilayer presented cross-linker are ordered membrane domains.

### 3.4 Membrane domain partitioning is not sufficient to describe CD45 and Lyn partitioning with respect to bilayer-engaged BCR clusters

In past work we found that CD45 exclusion from BCR clusters formed in response to crosslinking αIgM f(Ab)_1_ by soluble streptavidin could be attributed to CD45’s partitioning with membrane domains (Stone et al., 2017a). This conclusion was drawn because the truncated portion of CD45’s minimal membrane anchor sorted away from BCR clusters with the same magnitude as the full-length protein. In contrast, full-length Lyn kinase recruitment to BCR clusters was enhanced compared to that of its truncated minimal anchor portion, likely due to protein-protein interactions between full-length Lyn and proteins that reside in the BCR signalosome. These past findings are corroborated by images and cross-correlation functions in Fig 5A showing reconstructed images of CD45 and Lyn alongside αIgM labeled BCR clusters cross-linked via soluble streptavidin. Quantification over multiple images (Fig 5B) indicates that the subtle exclusion of CD45 from BCR clusters is similar in magnitude to that ob served for the Ld phase marker peptide TM. In contrast, Lyn colocalization with BCR clusters is more striking in reconstructed images and quantifications reveal that Lyn is more strongly recruited to BCR clusters than the Lo phase marker peptide PM. Representative images for PM and TM alongside IgM engaged with soluble cross-linker alongside average cross-correlation curves for BCR×TM and BCR×PM expressing cells are shown in Supplementary Figure 6.

**Figure 5:**
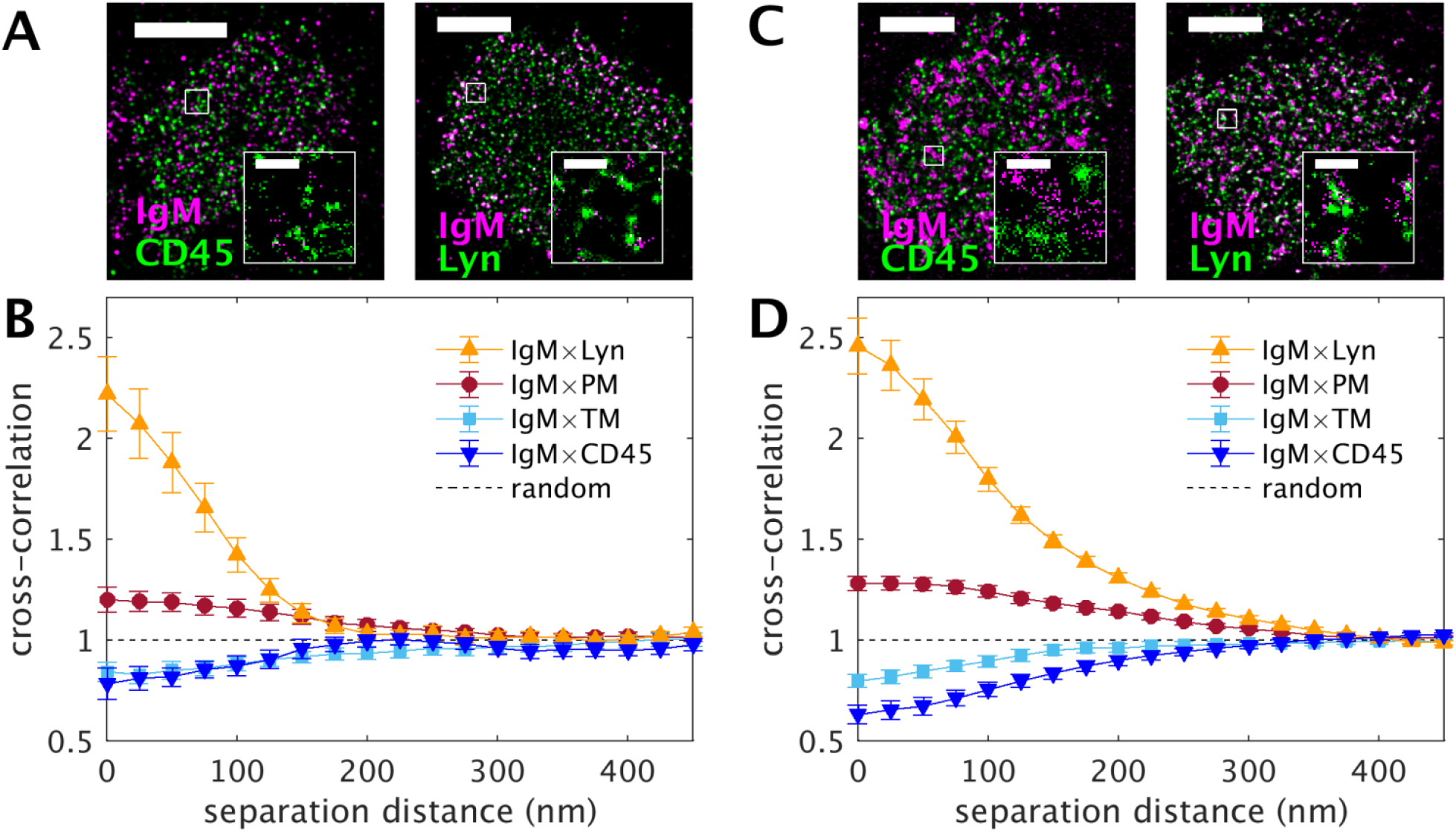
Full length regulatory proteins sort with respect to BCR clusters. (A) Representative images of CD45 and Lyn alongside BCR clusters formed by artificially crosslinking biotin-f(Ab)1 αIgM with soluble streptavidin. (B) Average cross-correlations between CD45 and Lyn with respect to IgM-BCR clusters tabulated from 20 and 22 cells respectively. These curves are plotted alongside the IgM×PM and IgM×TM curves for this same stimulation condition from Supplementary Figure 6. (C) Representative reconstructed super-resolution images of CD45 and Lyn alongside IgM-BCR clusters formed by artificially crosslinking biotin-f(Ab)_1_ αIgM with streptavidin presented on a planar lipid bilayer. (D) Average cross-correlations between CD45 and Lyn with respect to IgM-BCR clusters tabulated from 28 and32 cells respectively. These curves are plotted alongside the IgM×PM and IgM×TM curves for this same stimulation condition from Fig 4. Some data used to generate part B of this figure was published previously (Stone et al., 2017a). Scale bars in parts A and C are 5μm in main images and 500nm in insets.

We next explored the sorting of CD45 and Lyn in in BCR clusters formed in response to cross-linking with streptavidin presented on a supported membrane. Representative images and quantifications shown in Fig 5 C-D indicate that CD45 is more robustly excluded from BCR clusters than the minimal peptide TM, suggesting that coupling to membrane domains only provides a subset of the energy required to describe the organization of this protein. This observation is consistent with past work showing that proteins can be excluded from areas of close contact between the membrane and an opposing surface, sterically excluding proteins containing large ectodomains such as CD45 (Zhu et al., 2008;Harwood and Batista, 2009). For Lyn, results with surface presented streptavidin cross-linkers indicate that Lyn is more strongly recruited to BCR clusters than the ordered marker peptide PM, similar to observations with soluble streptavidin shown in Fig 5B. This again indicates that additional interactions contribute to Lyn’s recruitment to BCR clusters. Lyn was transiently transfected in these measurements while endogenous CD45 was antibody labeled. In all cases, we detect no correlation between the amplitude of g(r) from single cells and the expression level of these proteins (Supplemental Figure 7).

### 3.5 BCR clusters engaged with surface presented cross-linkers are highly tyrosine phosphorylated

Lyn is a kinase involved in phosphorylating the ITAM domains of BCR and other components of the signaling complex at tyrosine residues, whereas CD45 is one of several phosphatases that remove tyrosine phosphorylation (Katagiri et al., 1995;Katagiri et al., 1999;Cordoba et al., 2013). Our observations of enhanced sorting of the regulatory proteins CD45 and Lyn in cells stimulated with bilayer presented streptavidin led us to speculate that BCR clusters in these cells are more highly tyrosine phosphorylated than those found in clusters engaged with soluble streptavidin. To test this hypothesis, we visualized BCR alongside an antibody against total phosphotyrosine (pTyr) by diffraction limited and super-resolution localization microscopy and results are summarized in Fig 6.

**Figure 6:**
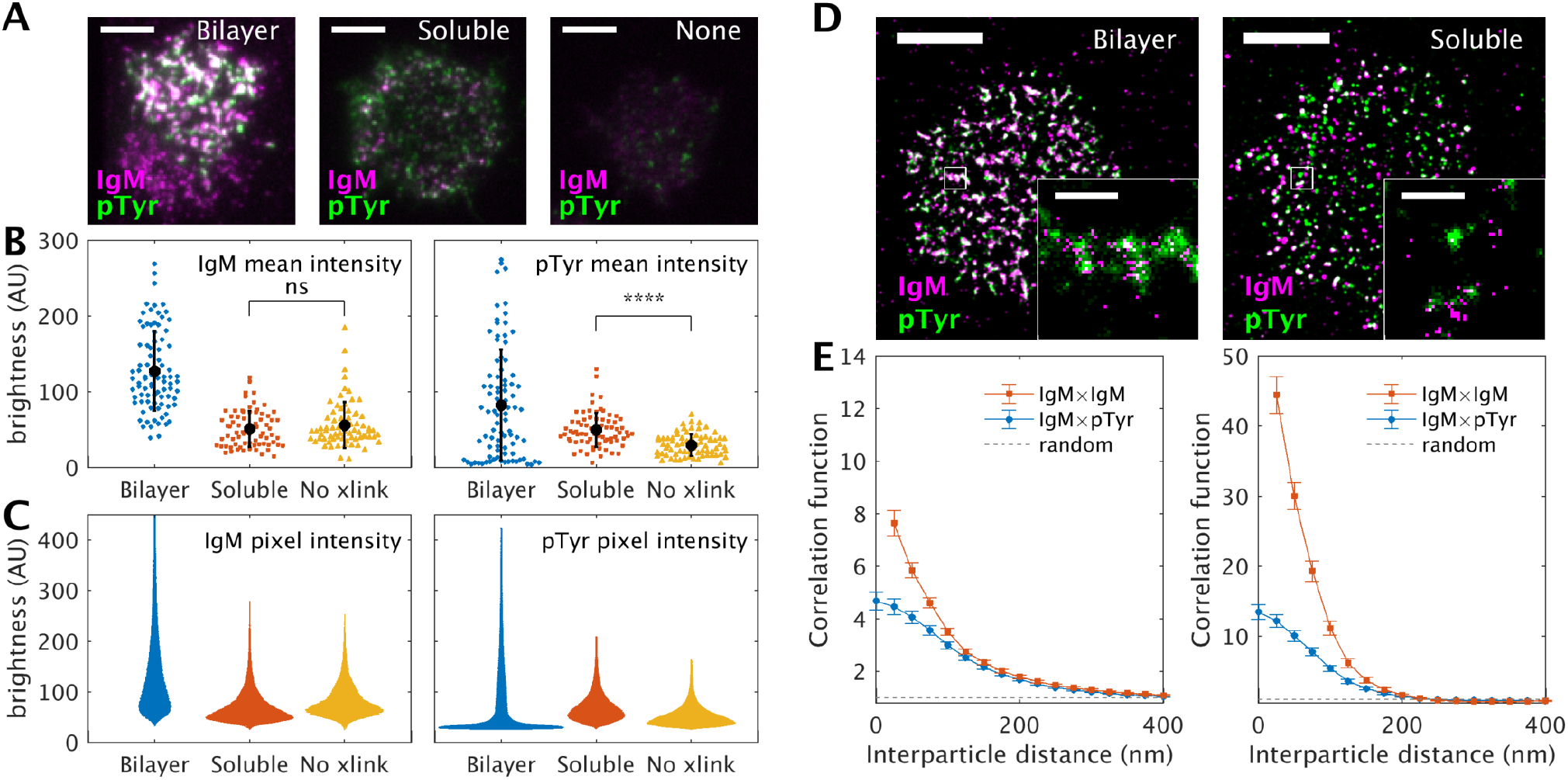
Tyrosine phosphorylation (pTyr) of artificially cross-linked IgM-BCR clusters. (A) Diffraction limited images of f(Ab)_1_ αIgM and total antibody stained pTyr in CH27 cells. Cells were imaged in TIR under the same illumination conditions and are displayed with the same brightness levels to enable comparison between conditions. (B) Quantification showing the average brightness across the cell membrane of over <100 individual cells for the three conditions shown in each color channel. (C) Quantification showing the distribution of pixel intensities for the same images quantified in part B. In this representation, the width of the bar indicates the fraction of pixels with that brightness level. (D) Representative reconstructed super-resolution images showing IgM and pTyr spatial distributions in CH27 cells engaged with bilayer presented or soluble cross-linker. Scale bars are 5μm in main images and 500nm in insets. (E) Average cross-correlation functions between IgM and pTyr contrasted with average auto-correlations for IgM for the two stimulation conditions shown in part D. Curves are tabulated from 33 cells with the bilayer cross-linker and 38 cells with the soluble cross-linker.

Figure 6A shows representative diffraction limited TIR images of cross-linked BCR (αIgM) and pTyr in cells stimulated with the surface presented or soluble artificial ligand streptavidin, or in control cells lacking exposure to either artificial cross-linkers or natural ligands. From the images it is clear that αIgM BCR density at the basal surface is increased in cells stimulated with bilayer-presented streptavidin. This is likely because biotin-f(Ab)_1_ αIgM labeled BCRs present on the apical surface of the cell either diffuse or are trafficked to the basal surface where the biotin engages streptavidin, ultimately concentrating labeled BCR. Absolute pTyr levels are also higher in cells engaged with bilayer-presented streptavidin compared to soluble streptavidin. Qualitative observations are quantified in Figure 6B, which report the average brightness of fluorescent probes over the cell-bilayer contact area in each color detected in more than 50 individual cells per condition. This quantification verifies that the average intensity of both αIgM and pTyr is enhanced in cells engaged with an artificial ligand-presenting supported lipid bilayer. Fig 6C shows histograms of pixel intensities reflective of the local density of components for the same cells quantified for Fig 6B. These histograms indicate that most αIgM labeled BCR is localized to high density clusters when engaged with bilayer-presented streptavidin, but clusters are small and sparse in cells stimulated with soluble streptavidin. Pixel intensities for pTyr staining indicate a more general enhancement of signal in cells stimulated with soluble streptavidin, and a bimodal distribution for cells stimulated with bilayer presented streptavidin. pTyr is highly enriched in some regions, likely coinciding with BCR clusters.

Surprisingly, a large number of pixels in surface engaged cells have very low pTyr intensities, below that of unstimulated cells. In addition, we sometimes observe that pTyr distributions appear to be polarized within cells stimulated with bilayer-presented streptavidin, with some areas of the cell-bilayer contact area staining highly for pTyr while others exhibit low pTyr staining, even when BCR clusters are present. Cells observed to be polarized in this way are often aligned with the direction of fluid flow in the chamber, suggesting that the mode of attachment to the bilayer may play some role in this phenomenon. An example of one of these cells is shown in the left panel of Fig 6A, which shows high pTyr staining on the top portion of the cell highly colocalized with IgM (white regions) and low pTyr staining on the bottom even though IgM clusters are still present (magenta regions). Numerous past studies have detailed the polarization or capping that occurs in response to soluble natural ligands or cross-linkers, or the formation of synapses in response to surface-presented antigens. Typically these processes are observed when cells are stimulated at 37°C or when cells are examined after longer stimulation times than investigated here (Batista et al., 2001; Carrasco et al., 2004; Fleire et al., 2006; Ketchum et al., 2014; Lee et al., 2017; Liu et al., 2010; Sohn et al., 2008a; Sohn et al., 2008b).

The co-localization of αIgM and pTyr signals was quantified from super-resolution images of samples engaged with bilayer-presented and soluble streptavidin cross-linkers, and representative images are shown in Fig 6D. In both cases, pTyr staining is visibly co-localized with BCR clusters. This observation is quantified by the cross-correlation functions in Figure 6E that produce curves with very high cross-correlation amplitudes. To assess how stringently the pTyr signal follows that of BCR, we contrast the pTyr αIgM cross-correlation to the αIgM autocorrelation (Fig 6E). For cells activated with bi-layer-presented streptavidin, cross-correlation and auto-correlation curves have similar amplitudes and shapes, indicating that pTyr signals largely sample the BCR distribution in these cells. In contrast, cells activated with soluble streptavidin produce cross-correlation and autocorrelation curves that have very different amplitudes, indicating more pTyr staining occurs away from BCR clusters under this stimulation condition. These observations are in good agreement with the diffraction limited findings of Fig 6C which show a global increase in pTyr staining across cells activated with the soluble cross-linker streptavidin and a more localized pTyr signal in cells activated with the bilayer-presented cross-linker streptavidin. We note that higher cross-correlation amplitudes for BCR engaged with soluble crosslinker do not indicate that these cells are more robustly tyrosine phosphorylated than bilayer cross-linked cells. Instead, this reflects that IgM domains engaged with soluble cross-linker occupy a smaller fraction of the cell surface under this stimulation condition, as explained more thoroughly in the discussion section. Taken together, these results indicate that BCR clusters are more strongly activated by surface-presented artificial cross-linking ligands and that this method of B cell stimulation strongly localizes pTyr activity to BCR clusters.

### 3.6 Membrane domains and phosphatase exclusion synergize collective receptor activation in a minimal model of early BCR signaling

In past work, we assembled a minimal model that produced collective receptor activation upon clustering within a heterogeneous membrane (Stone et al., 2017a). Within this model, receptor clustering leads to the stabilization of an extended ordered membrane domain that enriches kinase and depletes phosphatase based solely on their differential partitioning with membrane domains. Both receptors and kinase are represented as typical components that prefer ordered domains, phosphatases are represented as a typical component that prefers disordered domains. All proteins are embedded in a supercritical membrane that contains fluctuating domains and diffusive dynamics. Receptors are activated with a low probability when they come into contact with a kinase and are deactivated with high probability when they come into contact with a phosphatase. Additionally, activated receptors have kinase activity to mimic kinase binding to phosphorylated receptors, inserting a positive feedback loop that enhances the signaling output. An equal number of phosphatase and kinase components are included, and parameters are chosen such that the receptor activation state is low when receptors are not clustered in all models.

In the current study, we found that membrane coupling alone is not sufficient to describe phosphatase exclusion from BCR clusters formed through interactions with the bilayer presented cross-linker streptavidin. To account for this, we have updated the model to include an additional repulsive interaction sensed only by the phosphatase within the receptor cluster to mimic the potential experienced by proteins with bulky extracellular domains in regions of tight contact between the B cell plasma membrane and an antigen presenting cell. This additional repulsive interaction acts to reduce the local concentration of phosphatase within a localized membrane area, and here we implemented a large potential to enforce complete exclusion of phosphatase within receptor clusters. In sum, four minimal models were interrogated. One contains no interactions other than the potential used to cluster receptors, while the remaining models probe systems with domains and with phosphatase exclusion individually and in combination. Representative simulation snapshots are shown in Fig 7A and several model outcomes are shown in Figs 7B-D.

**Figure 7:**
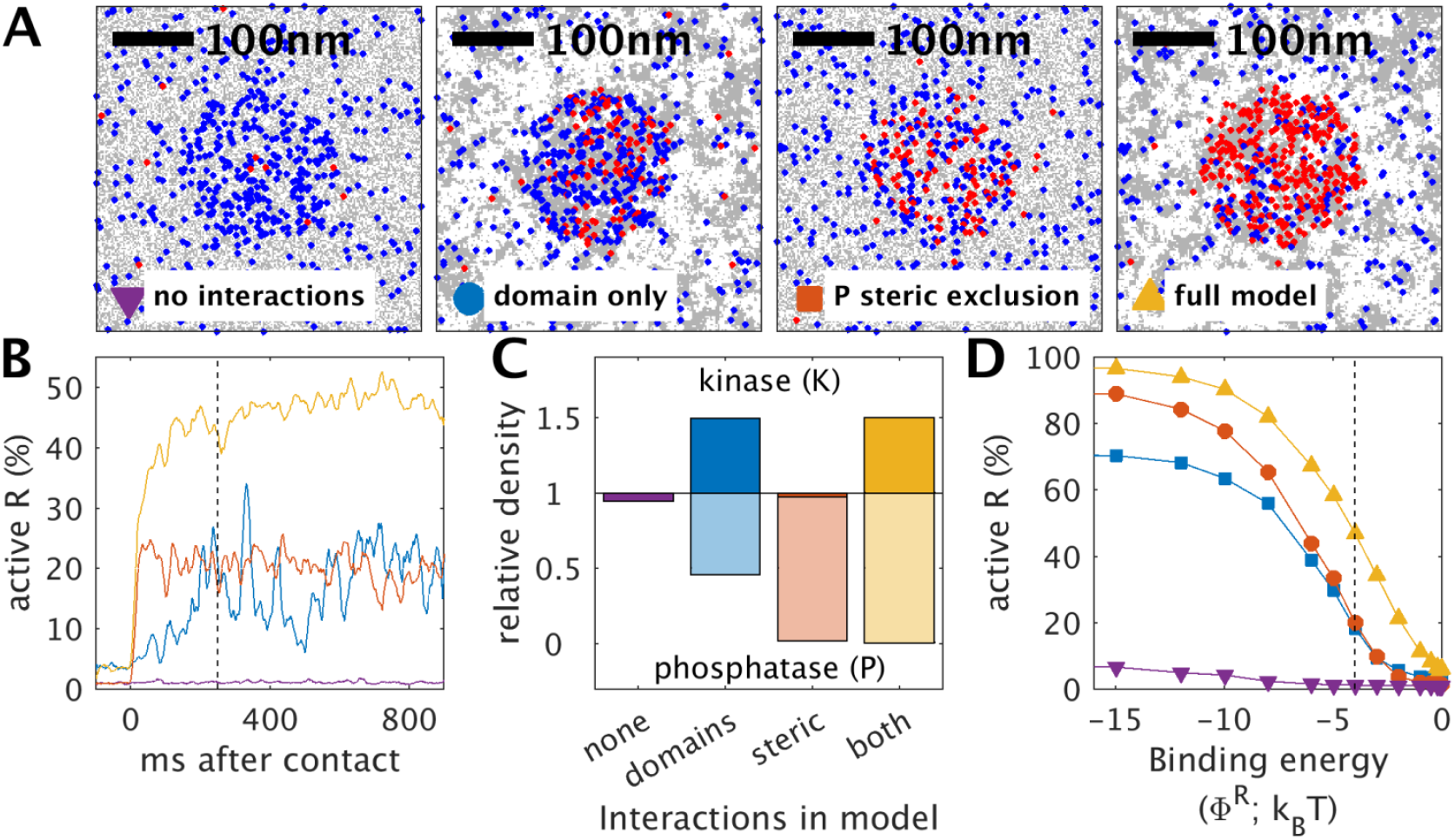
Simulations of a minimal model of receptor activation upon clustering. (A) Snapshots showing receptors and membrane domains taken from simulations of the 4 minimal models described in the main text. Receptors colored red are in an active state and receptors colored blue are in an inactive state. Receptors are clustered within a circle within the simulation frame through an applied field felt only by receptors and snapshots are generated from simulation results 250ms after the application of this field. (B) Traces showing the average number of active receptors within the simulation frame as a function of time for the 4 models investigated. The dashed vertical line represents the time corresponding to the snapshots shown in A. (C) The relative concentration of kinase and phosphatase within the receptor cluster for each of the 4 models investigated. Concentration is normalized to the concentration of each component over the whole simulation frame. (D) Average receptor activation as a function of ligand affinity obtained by tuning the strength of the receptor clustering field Φ^R^. Values are obtained by averaging values between 200ms and 1s after the application of the receptor clustering field. The dashed line indicates the receptor field strength corresponding to the snapshots shown in part A.

We find that receptors collectively activate upon clustering in the three models including domains and/or steric exclusion of phosphatase, although the fraction of activated receptors varies between models. This is seen in the representative traces showing the percentage of receptors in an activated state (Fig 7B) taken from the same simulations that produced representative snapshots (Fig 7A). In these examples, the full model has a large and sustained response, the phosphatase exclusion model has a sustained but reduced response, and the domain only model responds but the activation level varies more in time. The average activation levels can be understood by examining the average relative concentrations of kinase and phosphatase within domains shown in Fig 7C. In the domain only model, phosphatases can enter the receptor cluster where they can deactivate many receptors, leading to the reduction in average activation and the increased variability in receptor activation over time. Receptors cannot be inactivated by phosphatase within receptor clusters in the models with direct phosphatase exclusion, leading to larger overall activation levels and less variability within time. The difference in amplitude between the full model and the model with only phosphatase exclusion can be attributed to the local kinase concentrations in these models. Without domains, the local kinase concentration mimics that of the membrane as a whole, while domains facilitate the recruitment of kinase to receptor clusters, enhancing receptor activation. In addition, the number of receptors found in clusters is generally larger for the same clustering potential in models with domains. This is because coupling to domains makes receptor recruitment more cooperative, with receptors feeling an additional effective potential to localize with the other ordered components within this membrane region.

Figure 7D shows how receptor activation varies with affinity of the receptor-ligand binding association, which is tuned by varying the strength of the receptor clustering field in the models (Φ^R^). By this measure, the domain only and phosphatase exclusion models yield roughly the same sensitivity but vary in their maximum response under conditions of complete receptor clustering. To some extent, this agreement is due to parameters chosen for these examples but is meant to demonstrate that collective receptor activation upon clustering can occur via each of these mechanisms individually. Both sensitivity and the maximum stimulation capacity are enhanced in the model that combines membrane domains and steric phosphatase exclusion, demonstrating the potential synergy between these two distinct activation mechanisms. These simulations are in agreement with past studies which show that surface-based stimulation of B cells enhances their response (Tew et al., 1990;Wykes et al., 1998;Kosco-Vilbois, 2003;Carrasco and Batista, 2006b;Fleire et al., 2006).

## 4 DISCUSSION

### 4.1 BCR clusters engaged with a bilayer-presented natural ligand or artificial cross-linker are larger and contain more receptors than those engaged with an artificial soluble cross-linker

Past work has demonstrated that BCR clustering is enhanced when receptors are engaged with surface-bound antigens via the participation of specific oligomerization motifs (Tolar et al., 2009;Tolar and Pierce, 2010). Engaging BCR through mobile antigen on a fluid lipid bilayer has been shown to further enhance B cell signaling likely because active processes within cells can more easily gather BCR-antigen complexes into large domains (Ketchum et al., 2014;Lee et al., 2017). In the current work, we observe that receptor clusters are larger and contain more receptors when cells engage with bilayer-presented natural ligands or artificial cross-linkers than when BCR is clustered with artificial soluble cross-linkers (Fig 6B,C and Supplementary Figure 2A), likely due to these previously described mechanisms.

It is also possible that engagement of natural ligands acts as an additional contribution to the enhanced BCR clustering in the bilayer-engaged cells investigated in the current work when cells are stimulated with bilayer-bound artificial cross-linkers. For technical reasons, PC-containing bilayers were used even when presenting the artificial cross-linker streptavidin to biotin-f(Ab)_1_-αIgM labeled cells, and it is likely that some BCRs engage with PC lipids either instead of or in addition to engagement with the streptavidin cross-linker. We found that biotin-f(Ab)_1_-αIgM labeled CH27 cells do not adhere strongly to PC-containing bilayers that lack streptavidin, so it is possible that f(Ab)_1_-αIgM binding to receptors interferes with receptor binding to this natural ligand to some extent. Regardless of the mechanisms that give rise to the larger BCR clusters formed via bilayer stimulation, these structures provide a robust platform for investigating the sorting of plasma membrane proteins and peptides via superresolution fluorescence localization microscopy. BCR clusters in bilayer-engaged cell membranes have characteristic length-scales that exceed 100nm on average (Supplementary Figure 2A), which are well suited for imaging by this method that typically resolves fluorophores with 20-30nm lateral resolution.

### 4.2 BCR clusters engaged with a bilayer presented natural ligand or artificial cross-linker both stabilize robust ordered phase-like domains

This study quantifies the membrane composition proximal to BCR clusters established through interactions with a natural ligand or artificial cross-linker presented on a planar lipid bilayer using multicolor super-resolution fluorescence localization microscopy. We use lipids with PC headgroups as a natural monovalent ligand to stimulate CH27 B cells, and bilayer presented streptavidin as an artificial crosslinker to cluster αIgM f(Ab)_1_ bound to BCR. As detailed in previous studies, we find BCR clusters are extended and often non-circular when cells are simulated with bilayer presented ligands or cross-linkers (Ketchum et al., 2014;Lee et al., 2017). These larger BCR clusters sort minimal peptide markers of Lo and Ld phases in isolated plasma membrane vesicles. BCR clusters are enriched in a peptide marker of the Lo phase (PM) and depleted of a peptide marker of the Ld phase (TM), in good agreement with past studies that characterized the BCR microenvironment as an ordered membrane domain (Cheng et al., 1999;Pierce, 2002;Sohn et al., 2006;Sohn et al., 2008;Stone et al., 2017a). Importantly, our consistent observations of peptide sorting with respect to BCR clusters engaged with natural and artificial activators supports the view that that domain stabilization is a general property of BCR clusters.

### 4.3 Large, bilayer engaged BCR clusters effectively partition phase-marking peptides

Unlike the smaller BCR clusters formed upon cross-linking with soluble streptavidin interrogated in our previous study, the peptide sorting with respect to larger BCR clusters formed upon engagement with bilayer conjugated cross-linker is subtly evident in in reconstructed super-resolved images (Fig 4 and Supplementary Fig 4,5). This is in part because larger BCR clusters are well sampled in space by peptides, enabling visual comparison of peptide density both within and outside of clusters without statistical analysis. These improved spatial statistics are also evident when examining quantifications across a population of cells with cross-correlation amplitudes showing reduced variability for conditions that produce large domains (e.g. compare Fig 4C to Supplementary Fig 7B).

The sorting of phase-marking peptides is also more apparent in images with larger BCR clusters because these larger clusters more efficiently partition these peptides. This is evident through consideration of the relative amplitudes of the cross-correlation functions for soluble vs. surface-stimulated BCR compared to the relative area fractions occupied by BCR clusters for soluble vs. surface-stimulated BCR. The amplitude of the cross-correlation function is the concentration of peptide within BCR clusters normalized by the average peptide concentration across the entire cell area, rather than the peptide concentration away from clusters. When BCR clusters occupy a significant fraction of the cell surface, this average density of peptide over the cell surface is larger than the average density of peptides outside of BCR clusters. A more intuitive quantity to compare to images is the partition coefficient, K_P_, which is often defined as the ratio of peptide concentrations within and outside of clusters, ρ_on_/ρ_off_, where ρ_on_ is the concentration of peptide within BCR clusters and ρ_off_ is the concentration of peptide away from BCR clusters.

In a simplified 2 state system, ρ_ave_ = Aρ_on_ + (1-A)ρ_off_, where A is the area fraction occupied by BCR clusters and ρ_ave_ is the average peptide concentration across the cell surface. The cross-correlation function at r<25nm is roughly g(r<25nm) = ρ_on_/ρ_ave_. Through algebraic manipulation: K_P_ = ρ_on_/ρ_off_ = (1-A)g/(1-Ag). A is close to zero and K_P_ is well approximated by g(r<25nm) when the area fraction occupied by BCR clusters is small. In Fig 4, g(r<25nm) is roughly 0.8 for BCR×TM and 1.3 for BCR×PM. If we assume that the area fraction taken up by clusters is A=1/4, then K_P_ becomes 0.75 for TM and 1.4 for PM. If A=1/2, then K_P_ becomes 0.67 for TM and 1.86 for PM. These K_P_ values are well within the range found for similar peptides with respect to phase separated domains in isolated GPMVs (Diaz-Rohrer et al., 2014) and stabilized phase-like domains in vesicles adhered to supported membranes (Zhao et al., 2013).

In this study, we have not attempted to measure the area fraction occupied by BCR clusters (A), although it varies significantly from cell to cell and over the stimulation modes investigated. These changes in A complicate a quantitative comparison of the amplitude of g(r) across stimulation conditions. Qualitatively, we can conclude that phase-marking peptides partition with higher contrast (K_P_ further from 1) when stimulated with surface presented ligands or cross-liners than with a soluble multivalent cross-linker. This may be because bilayer-stimulated cells tend to contain a higher density of receptors and other components of BCR signaling complexes that favor ordered domains, as shown in Figure 6.

Lastly, we consistently observe that the characteristic length-scale of correlations between BCR and PM is larger than the characteristic length-scale of anti-correlation between BCR and TM (Supplementary Figure 2). It is possible that this effect arises due to the three-dimensional nature of the membrane. Our current study does not explicitly take into account membrane topography therefore makes the assumption that membrane topography is minimal. If this assumption is violated, then cross-correlations between membrane-bound compounds would appear correlated over a length-scale set by this topography (Parmryd and Onfelt, 2013). It is more exciting to speculate that the difference in the range of peptide recruitment and exclusion from BCR clusters reflects hierarchical organization of BCR micro-clusters, which contain scaffolding elements, numerous signaling regulators, recycling machinery, and cytoskeleton. It is possible that the peptides used in this study as markers of ordered and disordered membrane domains also sense other aspects of micro-clusters, such as membrane curvature or electrostatics. Exploring these ideas is an active area of future research.

### 4.4 Membrane anchor sequences of CD45 and Lyn contribute to their localization with respect to BCR clusters

The results shown in Fig 5 demonstrate that both CD45 and Lyn sort with respect to BCR clusters and that their inherent coupling to membrane domains contributes to this localization. When f(Ab)_1_ αIgM labeled BCR is clustered with a soluble cross-linker, CD45 is excluded to the same magnitude as TM, a minimal peptide marker of disordered domains, indicating that CD45 partitioning with disordered domains is sufficient to give rise to the observed depletion. In contrast, CD45 is more excluded than TM from BCR clusters formed by the same cross-linker presented on a supported membrane. We find that CD45 is roughly twice as excluded from BCR clusters compared to TM (g(r<25nm) 0.80±0.03 for αIgM×TM vs. ~0.63±0.05 for αIgM×CD45). Correlation functions can be converted to the potential of mean force (PMF) through the relation PMF(r) = − k_B_T ln(g(r)) where the PMF is the effective potential between the visualized proteins required to give rise to the observed spatial co-distribution at equilibrium (Veatch et al., 2012). This analysis suggests that coupling to membrane domains provides roughly half of the energy required to describe CD45 exclusion (PMF(r<25nm) = 0.22±.04 k_B_T for αIgM×TM and 0.46±.07 k_B_T for αIgM×CD45), while the remaining energy must come from other types of interactions in this system.

CD45 and other phosphatases implicated in regulating BCR signaling, such as CD148 (Kishihara et al., 1993;Byth et al., 1996;Harwood and Batista, 2009), have large extracellular domains. Past studies have attributed phosphatase exclusion from immune receptor clusters engaged with surfaces to steric effects whereby these proteins are prevented from entering membrane regions that are in tight juxtaposition to an opposing membrane (Harwood and Batista, 2008;Zhu et al., 2008). Overall, our results are consistent with this model, often referred to as the kinetic segregation model (Davis and van der Merwe, 2006), which has been studied more extensively in the context of T cell and macrophage signaling (Mustelin et al., 1989;Kishihara et al., 1993;Cordoba et al., 2013;Razvag et al., 2018).

In general, receptors tend be inactive when the effective activity of the phosphatase outpaces that of the kinase, and will tend to activate when either kinase activity increases or phosphatase activity decreases. Past work using reconstituted elements of the T cell receptor system have demonstrated that this biochemical network is highly sensitive to small changes in in the concentrations of these key regulators (Hui and Vale, 2014). The simulations presented in Fig 7 mimic the key aspect of the kinetic segregation model, by including a repulsive interaction at sites of receptor engagement felt only by phosphatases. As described by others (Davis and van der Merwe, 2006;Choudhuri et al., 2009;Schmid et al., 2016;Junghans et al., 2018), we find this mechanism alone is capable of inducing receptor activation upon engagement, by lowering the concentration and therefore the activity of phosphatase. Our simulations additionally demonstrate that the sensitivity and dynamic range of this signaling system is enhanced when it is also coupled to membrane domains. Membrane domains facilitate this enhancement through several avenues. First, local kinase concentration is increased at sites of receptor engagement, enabling receptor activation with higher phosphatase concentrations or more robust activation at the same phosphatase levels. Domains also facilitate collective recruitment of receptors to sites of ligand engagement, effectively increasing the affinity of the ligand-receptor interaction. This occurs because ligand binding inherently compensates for the losses in the mixing entropy accrued by confining a receptor to the cluster in an otherwise uniform membrane. When membrane domains are present, the free energy cost to organizing receptors is lowered, therefore less binding energy is required to recruit the same number of receptors. Overall, combining these mechanisms emphasizes the versatility of this signaling system, which is designed to be reactive to a broad range of stimuli.

### 4.5 Tyrosine phosphorylation activity increases at sites of enhanced protein sorting

Lyn and CD45 are involved in regulating early stages of BCR tyrosine phosphorylation, as well as the pTyr state of scaffolding elements and other signaling regulators within the BCR activation pathway. As a consequence, enhanced spatial sorting of Lyn and CD45 in the case of surface presented crosslinker is expected to give rise to elevated pTyr levels in the vicinity of BCR clusters, consistent with the experimental observations summarized in Fig 6. Additionally, the modeling presented in Fig 7 is consistent with the finding that surface engaged receptors become more active. Our model does not explicitly incorporate the enhanced kinase recruitment observed in the experiment, but it occurs indirectly since activated receptors have kinase activity. Within the model, we envision that the majority receptor bound kinases represent soluble kinases such as Syk, but could also include some quantity of Lyn. Overall, we find that receptor activation is enhanced under conditions where phosphatases are maximally excluded from receptor clusters.

Interestingly, we also observed surprisingly low pTyr levels away from BCR clusters formed via interactions with bilayer-presented vs. soluble cross-linkers. This is seen in the analysis of diffraction limited images in Fig 6C which show a large number of pixels with pTyr intensity lower than the no cross-linker control. This is also seen in the cross-correlations between IgM and pTyr of Fig 6E, indicating that pTyr more closely follows BCR structures in bilayer-presented compared to soluble crosslinker. It is possible that this occurs because the phosphatase concentration away from BCR is elevated under the condition of bilayer-presented cross-linker. It is tempting to speculate that changes in membrane protein and lipid composition away from receptors themselves contribute to signaling outcomes.

### 4.6 Summary and concluding remarks

In this study, we show that BCR clusters sort minimal markers of ordered and disordered phases when engaged with a bilayer presented monovalent ligand or with a bilayer conjugated artificial cross-linker, supporting the conclusion that these clusters are ordered, phase-like domains. Moreover, peptide sorting is more robust in cells stimulated with bilayer presented cross-linkers than in cells stimulated with soluble cross-linkers. Enhanced sorting of the regulatory proteins CD45 and Lyn is also observed in bilayer engaged cells. In particular, CD45 exclusion from bilayer engaged BCR clusters can only be partly attributed to its coupling to membrane domains, indicating other mechanisms contribute to the organization of this protein. This is consistent with published work from other groups, attributing phosphatase exclusion to steric interactions between their large ectodomains and the opposing supported membrane. As expected, BCR clusters in cells stimulated with surface presented natural ligands or artificial cross-linkers are more highly tyrosine phosphorylated, and this result can be recapitulated in a minimal model of receptor activation upon clustering that includes a repulsive interaction for phosphatases at sites of receptor clustering. Simulations combining phosphatase exclusion with coupling to membrane domains acts to enhance the sensitivity and dynamic range of this signaling system. Overall, this work supports the concept that the BCR signaling system is versatile, and draws from a variety of different protein-protein and protein-lipid interactions to tune sensitivity to a broad range of stimuli.

## 5 Author contributions

MN, Sample preparation, data acquisition and analysis, writing of the original draft and editing; KW, Sample preparation and data acquisition; SV, Conceptualization, data acquisition and analysis, analytical method development, supervision, writing of the original draft, and editing.

## 6 Conflict of interest statement

The authors declare no conflicts of interest for the submitted work.

## 7 Acknowledgements and funding

We thank Sarah Shelby, Thomas Shaw, Ben Machta, and Julia Bourg for helpful conversations and assistance with some measurements, as well as Sarah Shelby for comments on the manuscript. Research was supported by the NIH (R01GM110052) and the NSF (MCB1552439).

## References

Baumgart, T., Hammond, A.T., Sengupta, P., Hess, S.T., Holowka, D.A., Baird, B.A., and Webb, W.W. (2007). Large-scale fluid/fluid phase separation of proteins and lipids in giant plasma membrane vesicles. Proc Natl Acad Sci U S A 104, 3165–3170.

Burns, M.C., Nouri, M., and Veatch, S.L. (2016). Spot size variation FCS in simulations of the 2D Ising model. J Phys D Appl Phys 49.

Byth, K.F., Conroy, L.A., Howlett, S., Smith, A.J., May, J., Alexander, D.R., and Holmes, N. (1996). CD45-null transgenic mice reveal a positive regulatory role for CD45 in early thymocyte development, in the selection of CD4+CD8+ thymocytes, and B cell maturation. J Exp Med 183, 1707–1718.

Carrasco, Y.R., and Batista, F.D. (2006a). B - cell activation by membrane - bound antigens is facilitated by the interaction of VLA -;4 with VCAM - 1. The EMBO journal 25, 889–899.

Carrasco, Y.R., and Batista, F.D. (2006b). B cell recognition of membrane-bound antigen: an exquisite way of sensing ligands. Curr Opin Immunol 18, 286–291.

Carrasco, Y.R., Fleire, S.J., Cameron, T., Dustin, M.L., and Batista, F.D. (2004). LFA-1/ICAM-1 interaction lowers the threshold of B cell activation by facilitating B cell adhesion and synapse formation. Immunity 20, 589–599.

Cheng, P.C., Dykstra, M.L., Mitchell, R.N., and Pierce, S.K. (1999). A role for lipid rafts in B cell antigen receptor signaling and antigen targeting. J Exp Med 190, 1549–1560.

Choudhuri, K., Parker, M., Milicic, A., Cole, D.K., Shaw, M.K., Sewell, A.K., Stewart-Jones, G., Dong, T., Gould, K.G., and Van Der Merwe, P.A. (2009). Peptide-major histocompatibility complex dimensions control proximal kinase-phosphatase balance during T cell activation. J Biol Chem 284, 26096–26105.

Cordoba, S.P., Choudhuri, K., Zhang, H., Bridge, M., Basat, A.B., Dustin, M.L., and Van Der Merwe, P.A. (2013). The large ectodomains of CD45 and CD148 regulate their segregation from and inhibition of ligated T-cell receptor. Blood 121, 4295–4302.

Dal Porto, J.M., Gauld, S.B., Merrell, K.T., Mills, D., Pugh-Bernard, A.E., and Cambier, J. (2004). B cell antigen receptor signaling 101. Mol Immunol 41, 599–613.

Davis, S.J., and Van Der Merwe, P.A. (2006). The kinetic-segregation model: TCR triggering and beyond. Nat Immunol 7, 803–809.

Depoil, D., Fleire, S., Treanor, B.L., Weber, M., Harwood, N.E., Marchbank, K.L., Tybulewicz, V.L., and Batista, F.D. (2008). CD19 is essential for B cell activation by promoting B cell receptor-antigen microcluster formation in response to membrane-bound ligand. Nat Immunol 9, 63–72.

Diaz-Rohrer, B.B., Levental, K.R., Simons, K., and Levental, I. (2014). Membrane raft association is a determinant of plasma membrane localization. Proc Natl Acad Sci U S A 111, 8500–8505.

Dilillo, D.J., Horikawa, M., and Tedder, T.F. (2011). B-lymphocyte effector functions in health and disease. Immunologic research 49, 281–292.

Fleire, S.J., Goldman, J.P., Carrasco, Y.R., Weber, M., Bray, D., and Batista, F.D. (2006). B cell ligand discrimination through a spreading and contraction response. Science 312, 738–741.

Gasparrini, F., Feest, C., Bruckbauer, A., Mattila, P.K., Muller, J., Nitschke, L., Bray, D., and Batista, F.D. (2016). Nanoscale organization and dynamics of the siglec CD22 cooperate with the cytoskeleton in restraining BCR signalling. EMBO J 35, 258–280.

Gold, M.R., and Reth, M.G. (2019). Antigen Receptor Function in the Context of the Nanoscale Organization of the B Cell Membrane. Annu Rev Immunol 37, 97–123.

Harwood, N.E., and Batista, F.D. (2008). New insights into the early molecular events underlying B cell activation. Immunity 28, 609–619.

Harwood, N.E., and Batista, F.D. (2009). Early events in B cell activation. Annual review of immunology 28, 185–210.

Haughton, G., Arnold, L.W., Bishop, G.A., and Mercolino, T.J. (1986). The CH series of murine B cell lymphomas: neoplastic analogues of Ly-1+ normal B cells. Immunol Rev 93, 35–51.

Heilemann, M., Van De Linde, S., Mukherjee, A., and Sauer, M. (2009). Super-resolution imaging with small organic fluorophores. Angew Chem Int Ed Engl 48, 6903–6908.

Howard, M., and Paul, W.E. (1983). Regulation of B-cell growth and differentiation by soluble factors. Annual review of immunology 1, 307–327.

Hui, E., and Vale, R.D. (2014). In vitro membrane reconstitution of the T-cell receptor proximal signaling network. Nat Struct Mol Biol 21, 133–142.

Junghans, V., Santos, A.M., Lui, Y., Davis, S.J., and Jonsson, P. (2018). Dimensions and Interactions of Large T-Cell Surface Proteins. Front Immunol 9, 2215.

Katagiri, T., Ogimoto, M., Hasegawa, K., Arimura, Y., Mitomo, K., Okada, M., Clark, M.R., Mizuno, K., and Yakura, H. (1999). CD45 negatively regulates lyn activity by dephosphorylating both positive and negative regulatory tyrosine residues in immature B cells. J Immunol 163, 1321–1326.

Katagiri, T., Ogimoto, M., Hasegawa, K., Mizuno, K., and Yakura, H. (1995). Selective regulation of Lyn tyrosine kinase by CD45 in immature B cells. J Biol Chem 270, 27987–27990.

Ketchum, C., Miller, H., Song, W., and Upadhyaya, A. (2014). Ligand mobility regulates B cell receptor clustering and signaling activation. Biophys J 106, 26–36.

Kishihara, K., Penninger, J., Wallace, V.A., Kundig, T.M., Kawai, K., Wakeham, A., Timms, E., Pfeffer, K., Ohashi, P.S., Thomas, M.L., and Et Al. (1993). Normal B lymphocyte development but impaired T cell maturation in CD45-exon6 protein tyrosine phosphatase-deficient mice. Cell 74, 143–156.

Klasener, K., Maity, P.C., Hobeika, E., Yang, J., and Reth, M. (2014). B cell activation involves nanoscale receptor reorganizations and inside-out signaling by Syk. Elife 3, e02069.

Kosco-Vilbois, M.H. (2003). Are follicular dendritic cells really good for nothing? Nature Reviews Immunology 3, 764.

Kurosaki, T., Shinohara, H., and Baba, Y. (2009). B cell signaling and fate decision. Annual review of immunology 28, 21–55.

Lebien, T.W., and Tedder, T.F. (2008). B lymphocytes: how they develop and function. Blood 112, 1570–1580.

Lee, J., Sengupta, P., Brzostowski, J., Lippincott-Schwartz, J., and Pierce, S.K. (2017). The nanoscale spatial organization of B-cell receptors on immunoglobulin M- and G-expressing human B-cells. Mol Biol Cell 28, 511–523.

Levental, I., Lingwood, D., Grzybek, M., Coskun, U., and Simons, K. (2010). Palmitoylation regulates raft affinity for the majority of integral raft proteins. Proc Natl Acad Sci U S A 107, 22050–22054.

Levental, K.R., and Levental, I. (2015). Isolation of giant plasma membrane vesicles for evaluation of plasma membrane structure and protein partitioning. Methods Mol Biol 1232, 65–77.

Lorent, J.H., Diaz-Rohrer, B., Lin, X., Spring, K., Gorfe, A.A., Levental, K.R., and Levental, I. (2017). Structural determinants and functional consequences of protein affinity for membrane rafts. Nat Commun 8, 1219.

Machta, B.B., Papanikolaou, S., Sethna, J.P., and Veatch, S.L. (2011). Minimal model of plasma membrane heterogeneity requires coupling cortical actin to criticality. Biophys J 100, 1668–1677.

Mercolino, T.J., Arnold, L.W., and Haughton, G. (1986). Phosphatidyl choline is recognized by a series of Ly-1+ murine B cell lymphomas specific for erythrocyte membranes. J Exp Med 163, 155–165.

Minguet, S., Klasener, K., Schaffer, A.M., Fiala, G.J., Osteso-Ibanez, T., Raute, K., Navarro-Lerida, I., Hartl, F.A., Seidl, M., Reth, M., and Del Pozo, M.A. (2017). Caveolin-1-dependent nanoscale organization of the BCR regulates B cell tolerance. Nat Immunol 18, 1150–1159.

Monroe, J.G., and Dorshkind, K. (2007). Fate decisions regulating bone marrow and peripheral B lymphocyte development. Advances in immunology 95, 1–50.

Mukherjee, S., Zhu, J., Zikherman, J., Parameswaran, R., Kadlecek, T.A., Wang, Q., Au-Yeung, B., Ploegh, H., Kuriyan, J., Das, J., and Weiss, A. (2013). Monovalent and multivalent ligation of the B cell receptor exhibit differential dependence upon Syk and Src family kinases. Sci Signal 6, ra1.

Mustelin, T., Coggeshall, K.M., and Altman, A. (1989). Rapid activation of the T-cell tyrosine protein kinase pp56lck by the CD45 phosphotyrosine phosphatase. Proc Natl Acad Sci U S A 86, 6302–6306.

Niiro, H., and Clark, E.A. (2002). Decision making in the immune system: regulation of B-cell fate by antigen-receptor signals. Nature Reviews Immunology 2, 945.

Parmryd, I., and Onfelt, B. (2013). Consequences of membrane topography. FEBS J 280, 2775–2784.

Pierce, S.K. (2002). Lipid rafts and B-cell activation. Nat Rev Immunol 2, 96–105.

Pierce, S.K., and Liu, W. (2010). The tipping points in the initiation of B cell signalling: how small changes make big differences. Nat Rev Immunol 10, 767–777.

Pyenta, P.S., Holowka, D., and Baird, B. (2001). Cross-correlation analysis of inner-leaflet-anchored green fluorescent protein co-redistributed with IgE receptors and outer leaflet lipid raft components. Biophys J 80, 2120–2132.

Razvag, Y., Neve-Oz, Y., Sajman, J., Reches, M., and Sherman, E. (2018). Nanoscale kinetic segregation of TCR and CD45 in engaged microvilli facilitates early T cell activation. Nat Commun 9, 732.

Rey-Suarez, I., Wheatley, B., Koo, P., Shu, Z., Mochrie, S., Song, W., Shroff, H., and Upadhyaya, A. (2019). N-WASP regulates the mobility of the B cell receptor and co-receptors during signaling activation. bioRxiv, 619627.

Schmid, E.M., Bakalar, M.H., Choudhuri, K., Weichsel, J., Ann, H.S., Geissler, P.L., Dustin, M.L., and Fletcher, D.A. (2016). Size-dependent protein segregation at membrane interfaces. Nature Physics 12, 704–+.

Sendecki, A.M., Poyton, M.F., Baxter, A.J., Yang, T., and Cremer, P.S. (2017). Supported Lipid Bilayers with Phosphatidylethanolamine as the Major Component. Langmuir 33, 13423–13429.

Sengupta, P., Jovanovic-Talisman, T., Skoko, D., Renz, M., Veatch, S.L., and Lippincott-Schwartz, J. (2011). Probing protein heterogeneity in the plasma membrane using PALM and pair correlation analysis. Nat Methods 8, 969–975.

Sohn, H.W., Tolar, P., Jin, T., and Pierce, S.K. (2006). Fluorescence resonance energy transfer in living cells reveals dynamic membrane changes in the initiation of B cell signaling. Proc Natl Acad Sci U S A 103, 8143–8148.

Sohn, H.W., Tolar, P., and Pierce, S.K. (2008). Membrane heterogeneities in the formation of B cell receptor-Lyn kinase microclusters and the immune synapse. J Cell Biol 182, 367–379.

Stone, M.B., Shelby, S.A., Núñez, M.F., Wisser, K., and Veatch, S.L. (2017a). Protein sorting by lipid phase-like domains supports emergent signaling function in B lymphocyte plasma membranes. Elife 6.

Stone, M.B., Shelby, S.A., and Veatch, S.L. (2017b). Super-Resolution Microscopy: Shedding Light on the Cellular Plasma Membrane. Chem Rev 117, 7457–7477.

Stone, M.B., and Veatch, S.L. (2014). Far-red organic fluorophores contain a fluorescent impurity. Chemphyschem 15, 2240–2246.

Stone, M.B., and Veatch, S.L. (2015). Steady-state cross-correlations for live two-colour super-resolution localization data sets. Nat Commun 6, 7347.

Tew, J.G., Kosco, M.H., Burton, G.F., and Szakal, A.K. (1990). Follicular dendritic cells as accessory cells. Immunological reviews 117, 185–211.

Tolar, P., Hanna, J., Krueger, P.D., and Pierce, S.K. (2009). The constant region of the membrane immunoglobulin mediates B cell-receptor clustering and signaling in response to membrane antigens. Immunity 30, 44–55.

Tolar, P., and Pierce, S.K. (2010). “A conformation-induced oligomerization model for B cell receptor microclustering and signaling,” in Immunological Synapse. Springer), 155–169.

Veatch, S.L., Cicuta, P., Sengupta, P., Honerkamp-Smith, A., Holowka, D., and Baird, B. (2008). Critical fluctuations in plasma membrane vesicles. ACS Chem Biol 3, 287–293.

Veatch, S.L., Machta, B.B., Shelby, S.A., Chiang, E.N., Holowka, D.A., and Baird, B.A. (2012). Correlation functions quantify super-resolution images and estimate apparent clustering due to over-counting. PLoS One 7, e31457.

Volkmann, C., Brings, N., Becker, M., Hobeika, E., Yang, J., and Reth, M. (2016). Molecular requirements of the B-cell antigen receptor for sensing monovalent antigens. EMBO J 35, 2371–2381.

Weber, I. (2003). Reflection interference contrast microscopy. Methods Enzymol 361, 34–47.

Wykes, M., Pombo, A., Jenkins, C., and Macpherson, G.G. (1998). Dendritic cells interact directly with naive B lymphocytes to transfer antigen and initiate class switching in a primary T-dependent response. J Immunol 161, 1313–1319.

Yamanashi, Y., Kakiuchi, T., Mizuguchi, J., Yamamoto, T., and Toyoshima, K. (1991). Association of B cell antigen receptor with protein tyrosine kinase Lyn. Science 251, 192–194.

Youinou, P. (2007). B cell conducts the lymphocyte orchestra. Journal of autoimmunity 28, 143–151.

Zhao, J., Wu, J., and Veatch, S.L. (2013). Adhesion stabilizes robust lipid heterogeneity in supercritical membranes at physiological temperature. Biophys J 104, 825–834.

Zhu, J.W., Brdicka, T., Katsumoto, T.R., Lin, J., and Weiss, A. (2008). Structurally distinct Phosphatases CD45 and CD148 both regulate B cell and macrophage immunoreceptor signaling. Immunity 28, 183–196.

