## Supplemental figures S1-S8 for "Membrane domains and phosphatase exclusion produce robust signaling responses in B cells engaged with natural ligands and artificial cross-linkers presented on lipid bilayers"

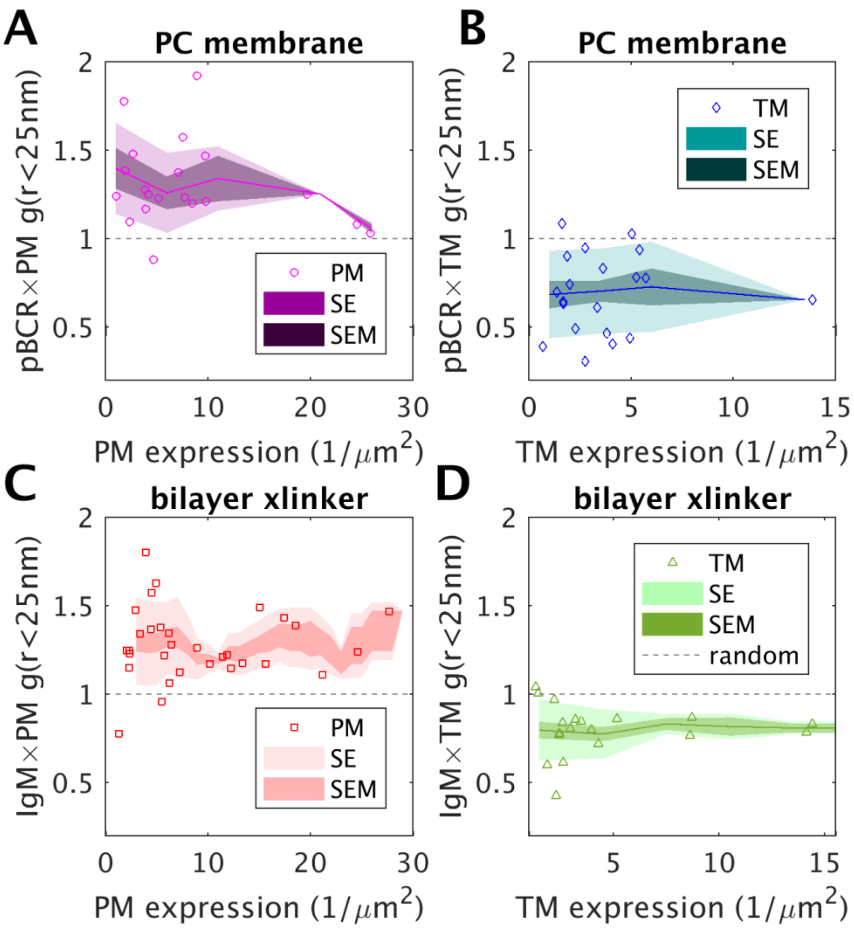


**Supplementary Figure 1: Cross-correlation amplitudes are independent of peptide expression levels.** (A,B) Points represent g(r<25nm) values of cross-correlation curves tabulated from individual cells for PM-mEos3.2 (A) and TM-mEos3.2 (B) in cells imaged after settling on a PC membrane. Expression levels are obtained by from the autocorrelation function as described previously (Veatch et al., 2012). The solid line shows a moving average of the points, the lighter shaded region shows the standard error and the darker shaded region shows the standard error of the mean. (C,D) Plots similar to the above but for biotin-f(Ab)$_1$-A655 labeled cells expressing PM-mEos3.2 (C) or TM-mEos3.2 (D) imaged after settling onto a membrane decorated with streptavidin cross-linkers.

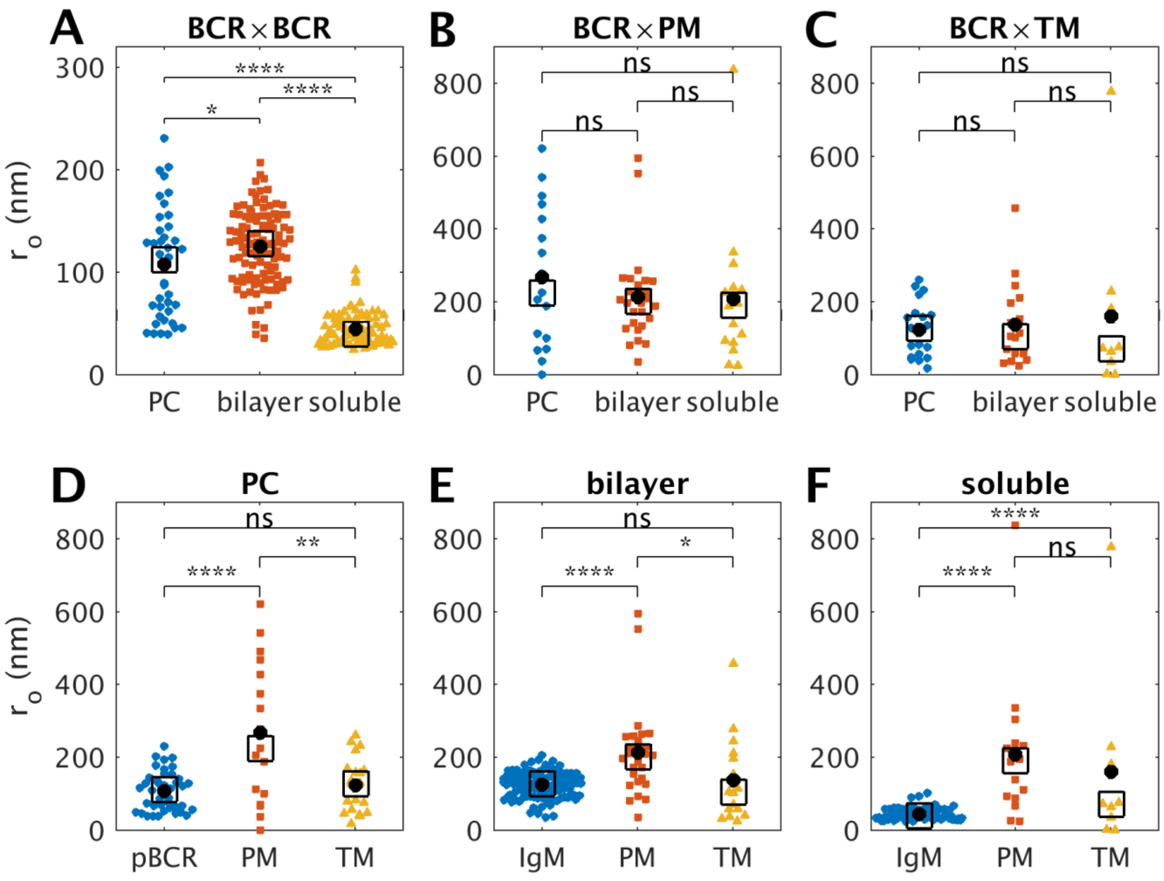


**Supplementary Figure 2: Characteristic length-scales of correlations across stimulation conditions.** (A-C) The characteristic length-scale ($r_o$) of correlations obtained by fitting BCR×BCR autocorrelation functions (A) and BCR×PM (B) and BCR×TM (C) cross-correlation functions to an exponential form, $g(r) = A\exp(-r/r_o) +$ offset for the stimulation conditions shown. pBCR is labeled for PC stimulation whereas IgM is labeled for stimulation via engagement with bilayer or surface presented cross-linkers. Colored points are obtained from single cells, the black circle indicates the mean and the square box is centered around the median. Outliers often occur when the overall amplitude of correlations in a cell is low, therefore the fitting to $r_o$ less reliable. More points are analyzed for BCR×BCR since cells labeled with PM, TM, Lyn, CD45, and pTyr were included. Significance was determined using a 2 sample t-test through the ttest2 command in Matlab. (D-F) The same results shown for parts A-C divided by stimulation type instead of probe pair.

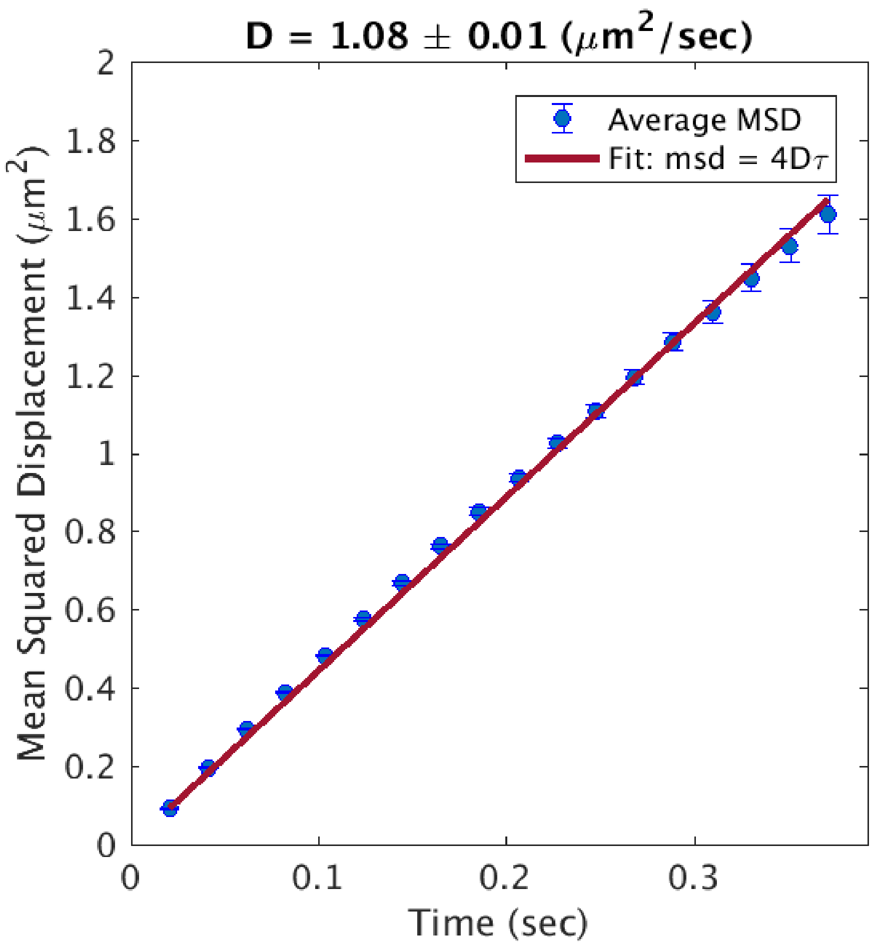


**Supplementary Figure 3: Streptavidin is mobile on supported lipid bilayers in our flow chambers.** Streptravidin-Alexa Fluor 647 fluorescence was tracked when loaded onto a supported lipid bilayer fused from large Unilamellar vesicles comprised of 99.5mol% DOPC and 0.5mol% biotinyl-cap-DOPE. The mean squared displacement of each single-molecule track was averaged by hundreds of localized probe tracks across 10 movies. Averaged streptavidin-AF647 MSD curve is presented with the standard error of the mean (blue). The diffusion coefficient of streptavidin was estimated by fitting the averaged MSD to a line (red) with slope = 4Dτ and a y-intercept fixed at 0. Streptavidin has a diffusion coefficient of 1.08 $\mu m^2$/sec, which is on the order of the diffusion coefficient of a mobile lipid in a bilayer and indicate that the antigen is mobile on the bilayers fused in our flow chambers.

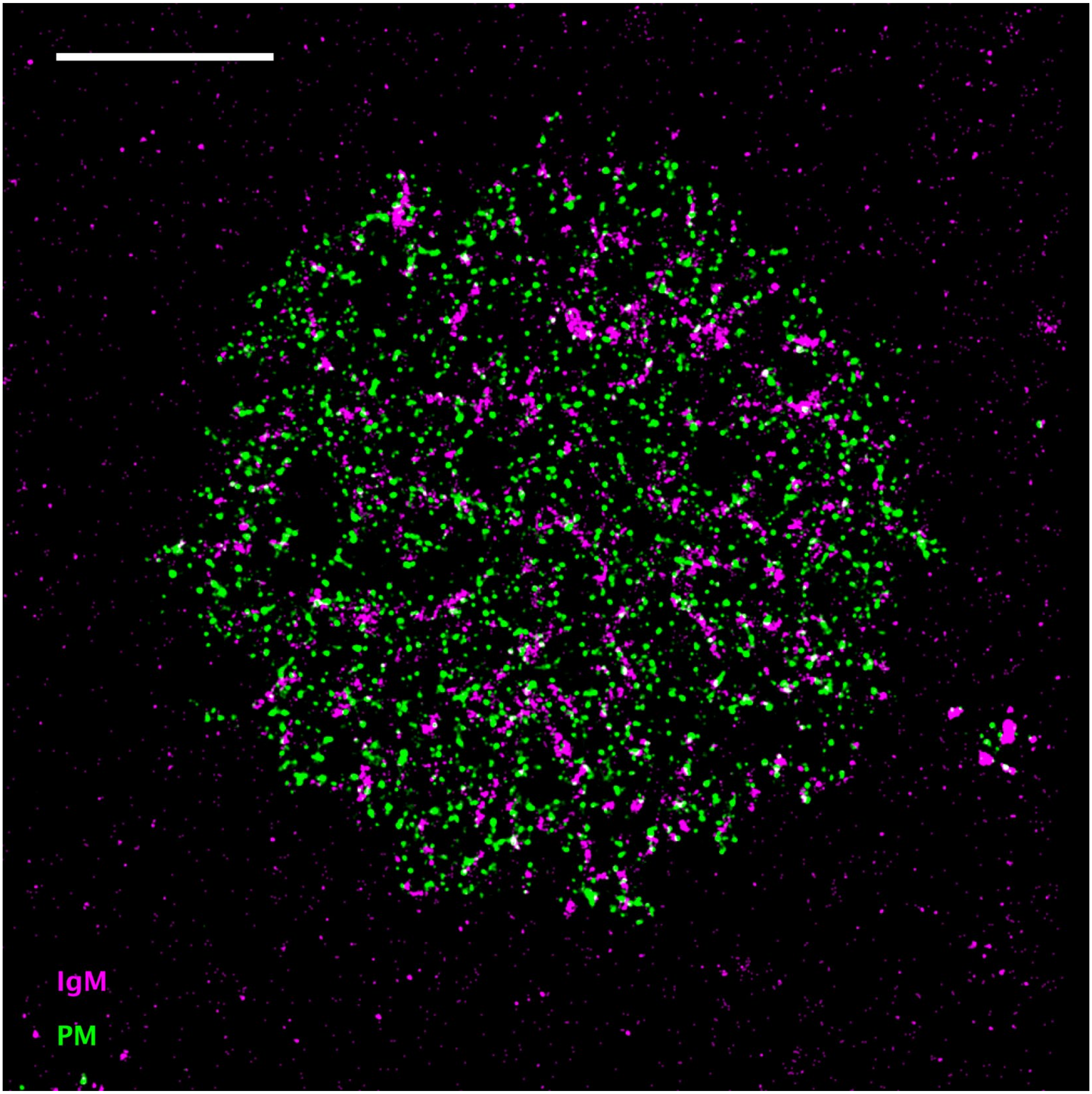


**Supplementary Figure 4: Reconstructed image of PM-mEos3.2 expressing cell engaged with PC bilayer presented streptavidin.** Image is reconstructed from grouped localizations with 25nm pixels and filtered with a 25nm Gaussian function. At a minimum, visual inspection indicates that green, mEos3.2 spots do not avoid magenta IgM clusters, and subtle enrichment may be apparent through close inspection. Compare with Supplementary Figure 5 which exhibits more exclusion. Scale bar is 5µm.

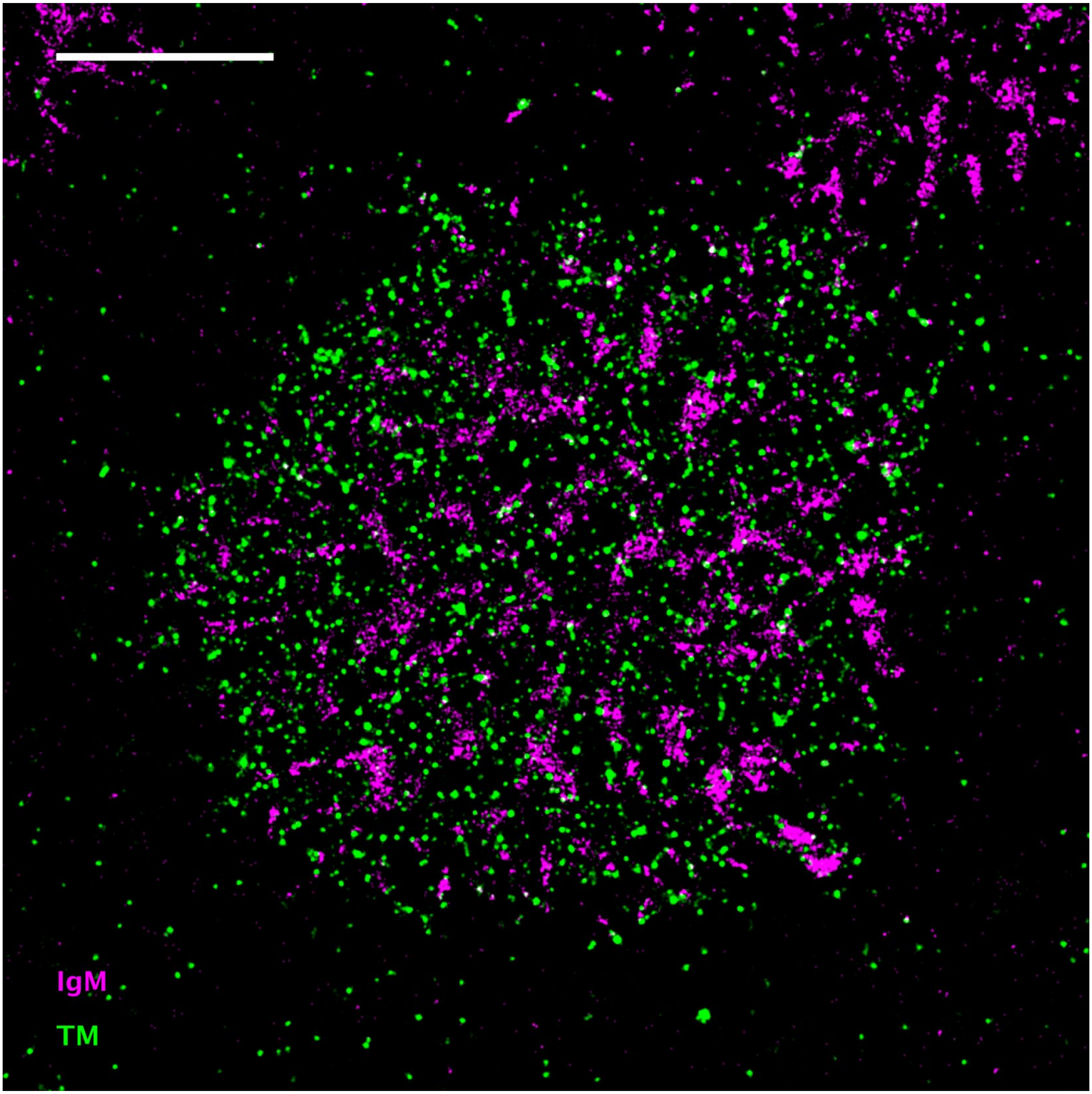


**Supplementary Figure 5: Reconstructed image of TM-mEos3.2 expressing cell engaged with PC bilayer presented streptavidin.** Image is reconstructed from grouped localizations with 25nm pixels and filtered with a 25nm Gaussian function. Visual inspection indicates that green, mEos3.2 avoid magenta IgM clusters. Compare with Supplementary Figure 5 which exhibits subtle enrichment. Scale bar is 5µm.

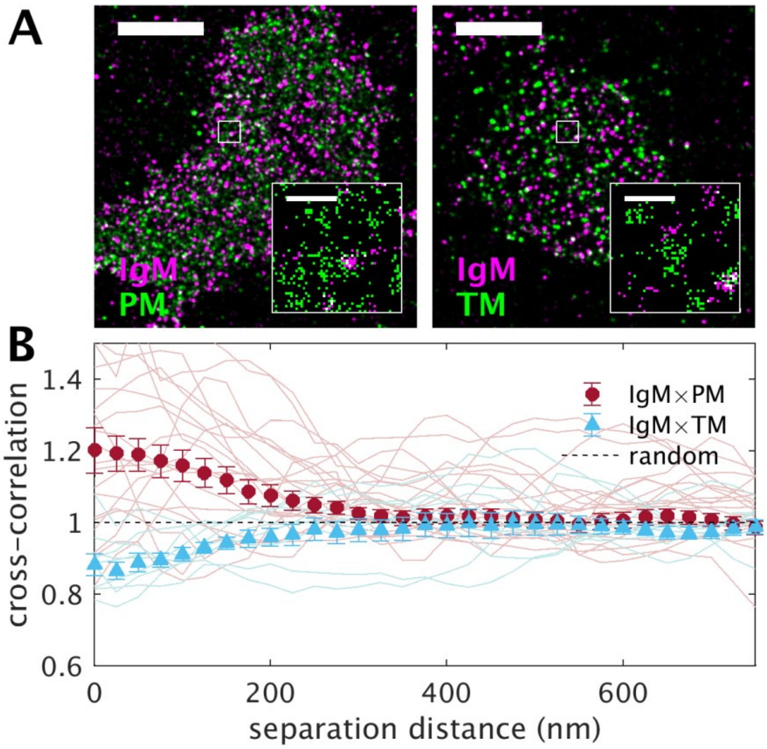


**Supplementary Figure 6: Phase-marking minimal peptides sort with respect to BCR clusters formed through engagement with the artificial cross-linker streptavidin administered in solution.**
(A) Representative reconstructed super-resolution images of BCR clusters, marked with biotinylated f(Ab)$_1$ αIgM conjugated to Atto655 and the minimal membrane anchored peptides PM (right) or TM (left) conjugated to mEos3.2. Scale bars are 5μm in main images and 500nm in insets. (B) Cross-correlation functions tabulated for 21 cells between IgM and PM or 10 cells IgM and TM. Curves from individual cells are shown as lighter color lines and average curves are shown as filled symbols with error bounds representing the standard error of the mean between cells. This figure includes some data published previously (Stone et al., 2017).

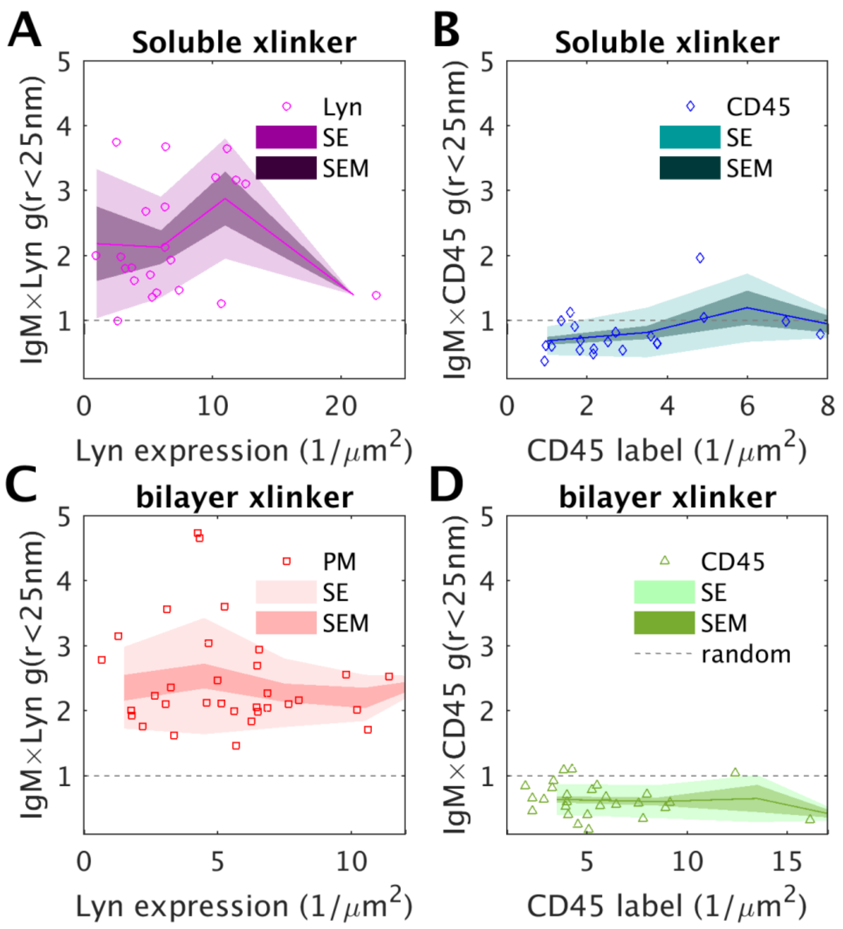


**Supplementary Figure 7: Cross-correlation amplitudes are independent of protein expression levels.** (A,B) Points represent g(r<25nm) values of cross-correlation curves tabulated from individual cells for Lyn-mEos3.2 (A) and antibody labeled endogenous CD45 (B) in cells imaged after having their IgM BCR cross-linked with soluble streptavidin. Expression levels are obtained by from the autocorrelation function as described previously (Veatch et al., 2012). The solid line shows a moving average of the points, the lighter shaded region shows the standard error and the darker shaded region shows the standard error of the mean. (C,D) Plots similar to the above but for cells engaged with streptavidin presented on a PC membrane or Lyn-mEos3.2 (C) and antibody labeled endogenous CD45 (D).

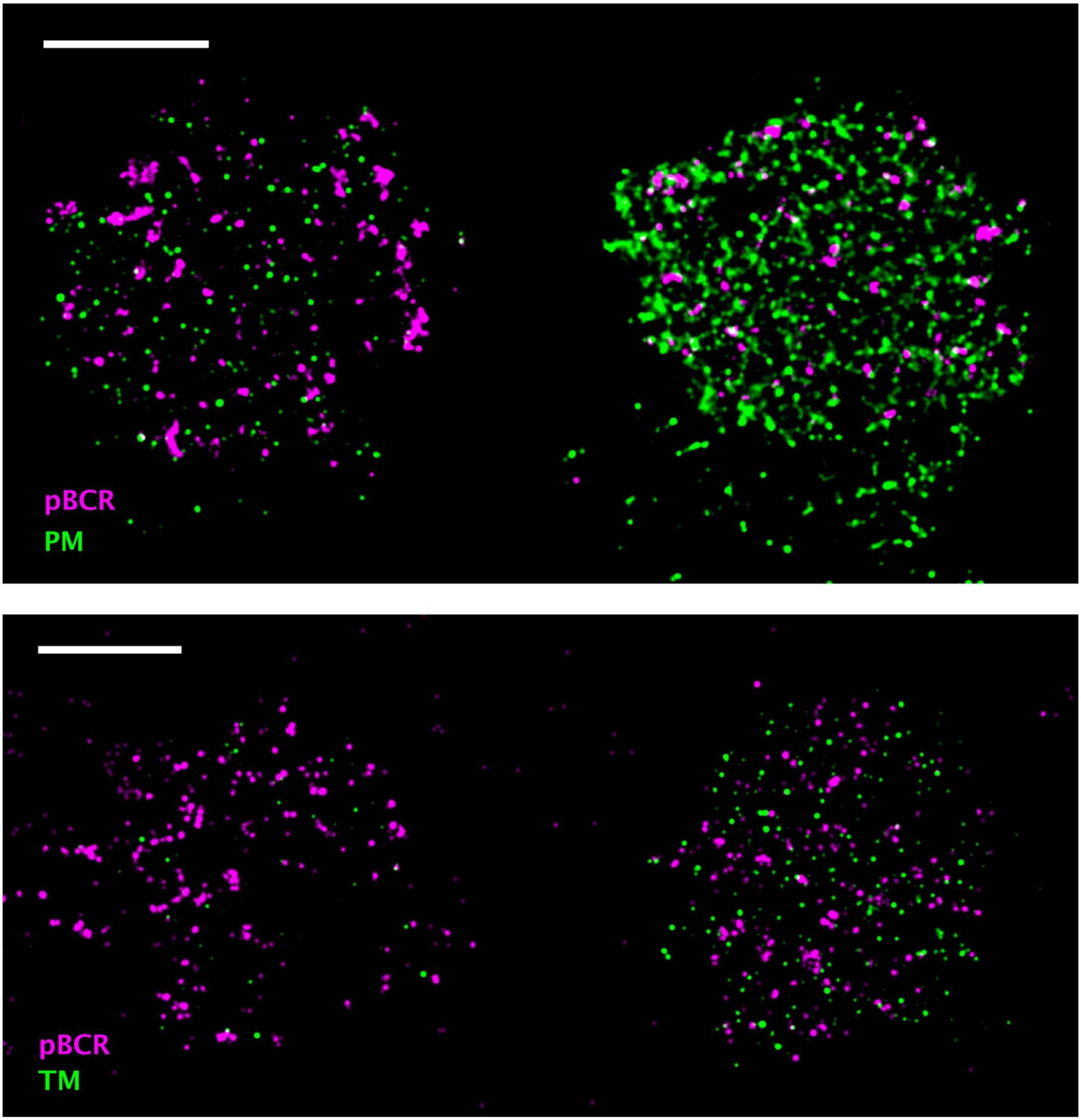


**Supplementary Figure 8: pBCR staining (magenta) of cells engaged with a PC membrane do not depend strongly on peptide expression level (green).** Images are reconstructed with 25nm pixels and are filtered with a 50nm Gaussian function.

### References

Stone, M.B., Shelby, S.A., Núñez, M.F., Wisser, K., and Veatch, S.L. (2017). Protein sorting by lipid phase-like domains supports emergent signaling function in B lymphocyte plasma membranes. *Elife* 6.

Veatch, S.L., Machta, B.B., Shelby, S.A., Chiang, E.N., Holowka, D.A., and Baird, B.A. (2012). Correlation functions quantify super-resolution images and estimate apparent clustering due to over-counting. *PLoS One* 7**,** e31457.
